# Phasic oxygen dynamics underlies fast choline-sensitive biosensor signals in the brain of behaving rodents

**DOI:** 10.1101/2020.08.05.237453

**Authors:** Ricardo M. Santos, Anton Sirota

## Abstract

Fast time-scale modulation of synaptic and cellular physiology by acetylcholine is critical for many cognitive functions, but direct local measurement of neuromodulator dynamics in freely-moving behaving animals is technically challenging. Recent *in vivo* brain measurements using choline oxidase (ChOx)-based electrochemical biosensors have reported surprising fast cholinergic transients associated with reward-related behavioral events. However, *in vivo* recordings with conventional ChOx biosensors could be biased by phasic local field potential and O_2_-evoked enzymatic responses. Here, we have developed a Tetrode-based Amperometric ChOx (TACO) sensor enabling minimally invasive artifact-free simultaneous measurement of cholinergic activity and O_2_. Strikingly, the TACO sensor revealed highly-correlated O_2_ and ChOx transients following spontaneous locomotion and sharp-wave/ripples fluctuations in the hippocampus of behaving rodents. Quantitative analysis of spontaneous activity, *in vivo* and *in vitro* exogenous O_2_ perturbations revealed a directional effect of O_2_ on ChOx phasic signals. Mathematical modeling of biosensors identified O_2_-evoked non-steadystate ChOx kinetics as a mechanism underlying artifactual biosensor phasic transients. This phasic O_2_-dependence of ChOx-based biosensor measurements confounds phasic cholinergic dynamics readout *in vivo,* challenging previously proposed ACh role in reward-related learning. The discovered mechanism and quantitative modeling is generalizable to any oxidase-based biosensor, entailing rigorous controls and new biosensor designs.

## Introduction

Acetylcholine (ACh) is an essential modulator of neuronal circuits engaged in high order cognitive operations. During aroused states, high extracellular ACh sets cortico-hippocampal circuits towards memory encoding by enhancing sensory processing, synaptic plasticity and neuronal network rhythmicity (Hasselmo and McGaughy; Marrosu et al., 1995; Steriade, 2004; Teles-Grilo Ruivo and Mellor, 2013). The latter is particularly relevant in the hippocampus, where ACh plays an important role in the processing of episodic and emotional information via modulation of theta oscillations (Buzsáki, 2002; Gu et al., 2017; Li et al., 2007; Mikulovic et al., 2018; Vandecasteele et al., 2014). Contrarily, low tonic ACh during non-REM sleep permits the occurrence of hippocampal sharp-wave/ripple complexes (SWRs), which are critical for memory consolidation (Buzsáki, 2015; Hasselmo and McGaughy, 2004; Norimoto et al., 2012; Vandecasteele et al., 2014).

The current theory on the functional role of ACh in cortical and hippocampal circuits has been mainly derived from brain-state-related correlations of tonic ACh levels with underlying network dynamics, or from strong manipulations of the cholinergic system (Gu et al., 2017; Hasselmo and McGaughy, 2004; Li et al., 2007; Marrosu et al., 1995; Norimoto et al., 2012; Vandecasteele et al., 2014). However, such crude analytic and experimental approaches cannot account for the spontaneous non-stationary interactions between ACh, behavior and neuronal network activity. Recently, fine time-scale measurements of cholinergic activity have provided new insights into these interactions. Fast cholinergic transients in cortical and hippocampal regions have been described in response to sensory sampling, unexpected events, negative reinforcements and reward-related behavior (Eggermann et al., 2014; Hangya et al., 2015; Howe et al., 2017; Lovett-Barron et al., 2014; Parikh et al., 2007; Teles-Grilo Ruivo et al., 2017). The latter have been captured using electrochemical biosensors in response to detection of cues to rewards and reward approach or retrieval in freely-moving rodents (Howe et al., 2017; Parikh et al., 2007; Teles-Grilo Ruivo et al., 2017). The temporally-precise alignment of phasic ACh signals to these events hints for a critical role of ACh on the formation of reward-related memories and on the guiding of learned reward-oriented actions.

The above-mentioned studies highlight the suitability of enzyme-based electrochemical biosensors to capture phasic release of neurotransmitters and neuromodulators(Chatard et al., 2018). Additionally, amperometric measurements pick-up currents generated by the local field potential (LFP) (Viggiano et al., 2012; Zhang et al., 2009), making these sensors ideal for studying the interplay between neuromodulatory tone and neuronal network dynamics. The most successful electrochemical ACh sensing strategy *in vivo* has relied on the Choline Oxidase (ChOx)-mediated measurement of extracellular choline (Ch), a product of ACh hydrolysis by acetylcholinesterase. The enzyme catalyzes Ch oxidation in the presence of O_2_, generating H_2_O_2_, which is oxidized on the electrode surface (Burmeister et al., 2003; Parikh et al., 2004, 2007; Teles-Grilo Ruivo et al., 2017; Zhang et al., 2010).

However, despite the apparent success of ChOx-biosensors, the factors that can confound their response *in vivo* at the fast time-scale in freely-moving animals, such as LFP-related artifacts and O_2_-evoked enzyme transients, have not been thoroughly addressed.

First, the approach that has been conventionally used to cancel LFP-related currents, based on the differential coating of the recording sites with matrices that contain (Ch-sensing sites) or lack ChOx (sentinel sites), is not optimal. Although differential measurements are essential to clean biosensor signals (Santos et al., 2015; Zhang et al., 2010), H_2_O_2_ diffusion from enzyme-coated to sentinel sites poses important constraints on the sensor design. The inter-site spacing required to avoid diffusional cross-talk leads to uncontrolled differences in the amplitude and phase of LFP across sites, compromising common-mode rejection.

Second, the strategies devised to reduce artifacts that directly generate electrochemical currents (chemical surface modifications or common-mode rejection) and are unrelated to enzymatic response to Ch, do not control for factors influencing ChOx activity. Given that O_2_ is a co-substrate of the enzyme, it is crucial to control whether physiological O_2_ variations can contribute to biosensor responses *in vivo* leading to distortion of true and detection of false positive Ch signals. Previous studies have only shown that O_2_ steady-state responses of ChOx-based biosensors *in vitro* follow apparent Michaelis-Menten saturation kinetics (Baker et al., 2017; Burmeister et al., 2003; Santos et al., 2015). The relatively narrow linear range of O_2_-dependent biosensor responses, as compared with estimates of average O_2_ levels in the brain, has motivated the assumption that ChOx-based biosensors are not affected by *in vivo* O_2_ dynamics (Baker et al., 2017; Burmeister et al., 2003). That might, however, oversimplify the effect of O_2_ on enzymatic activity since it widely varies across brain regions and experimental conditions (Lyons et al., 2016; Murr et al., 1994; Nair et al., 1987). Furthermore, previous literature has ignored possible non-steady-state (phasic) biosensor responses to O_2_, which might arise from local consumption and diffusion of enzyme substrates and reaction products in the sensor coating. These putative transient sensor responses to O_2_ are particularly relevant as they might temporally overlap with fast cholinergic transients. The full assessment of biosensors’ O_2_ dependence requires the characterization of tonic and phasic O_2_-evoked responses and the simultaneous *in vivo* measurement of immobilized ChOx activity (biosensor response) and O_2_ within the sensor substrate. Yet, when addressed, O_2_ levels in tissue have been measured using a separate electrode, often of different geometry and/or having a surface material or modification that differs from the Choline-sensing site (Baker et al., 2015; Dixon et al., 2002). This approach is therefore prone to bias from heterogeneous tissue O_2_ dynamics and differential kinetics of electrode responses to O_2_.

Here, we have circumvented the above-discussed limitations by designing a novel Tetrode-based Amperometric ChOx (TACO) sensor that provides differential responses to Ch, interferents and LFP across four bundled Pt wires. Importantly this multichannel configuration allows the unbiased measurement of the immobilized ChOx activity (COA) and O_2_ in the same brain spot by using a tetrode site to directly measure the latter. Experiments with TACO sensor in the hippocampus of freely-moving and head-fixed rodents highlight the relevance of our common-mode rejection strategy to provide highly-sensitive contamination-free COA and O_2_ signals. We found fast O_2_ and COA dynamics following locomotion bouts in active state and hippocampal SWRs during quiescence state. Remarkably, spontaneous O_2_ and COA transients correlated in amplitude and time regardless of the underlying behavioral or neurophysiological context, a phenomenon we reproduced via causal local or systemic manipulation of O_2_ dynamics. Our *in vivo* findings, together with a detailed *in vitro* characterization and mathematical modeling of O_2_-evoked biosensor signals reveal a major contribution of spontaneous O_2_ dynamics to phasic ChOx-based biosensor signals. Importantly, this O_2_ ChOx-confounding dynamics is associated with behaviorally and physiologically relevant events and warrants important implications for the interpretation of previous studies relying on ChOx sensors as well as for the design of the future enzyme-based sensors.

## Results

### Differential plating of TACO sensor discriminates choline from interferents

TACO sensor is built around Pt/Ir wire tetrode, providing a spatial density of recording sites that is ideal for common-mode rejection of LFP-related currents and neurochemical dynamics. At this spatial scale, diffusional crosstalk would preclude the use of sentinel sites by differential coatings, as done in conventional biosensors (Burmeister et al., 2003; Santos et al., 2015). Instead, we created *pseudo*sentinel and Ch-sensing sites by differentially plating the tetrode wires, modifying their electrocatalytic response towards H_2_O_2_. This step was followed by coating the tetrode surface with a common matrix containing ChOx entrapped in chitosan (Santos et al., 2015) (Figure 1A). As for the initial plating steps, all tetrode sites were first mildly plated with gold, which marginally increased the electrode surface area, as inferred from impedances at 1 kHz (419 ± 33 kΩ, n = 28 before vs. 370 ± 16 kΩ, n = 44 after gold plating). Despite the slight decrease in impedance, gold-plating significantly decreased the electrocatalysis of H_2_O_2_ reduction/oxidation at the metal surface. In enzyme-coated electrodes, gold-plated sites exhibited a nearly 5-fold smaller H_2_O_2_ sensitivity than an unplated Pt/Ir surface (Figure S1). Remarkably, following this first step, gold-plated sites could be rendered H_2_O_2_-sensitive upon mild platinization. In enzyme-coated electrodes, there was a nearly 10-fold difference in H_2_O_2_ oxidation currents between Au and Au/Pt sites at 0.4-0.6 V vs. Ag/AgCl (Figure 1B). The increase in Au sites’ response to H_2_O_2_ above +0.7 V is in agreement with the electrochemical behavior of a pure Au electrode and probably results from the formation of surface oxides (Burke and Nugent, 1997; O’Neill et al., 2004). Interestingly, the response of Au/Pt electrodes to H_2_O_2_ was higher than that of unplated electrodes (Figure S1), possibly reflecting an increase in the electrode surface area and/or an electrocatalytic effect caused by the deposition of nanostructured platinum over the gold surface (Burke, 1996; Domínguez-Domínguez et al., 2008). Even when using common-mode rejection, decreasing the magnitude of interferences in individual sites is desirable. Thus, after plating and coating the tetrode, we electropolymerized *m*-PD in two of the recording sites, in order to reduce responses to electroactive compounds larger than H_2_O_2_ (eg. ascorbate or dopamine) (Hascup et al., 2013; Santos et al., 2008). The final TACO sensor configuration consisted of all possible combinations of Pt and *m*-PD modifications of Au-plated recording sites (Figure 1A).

**Figure 1.**
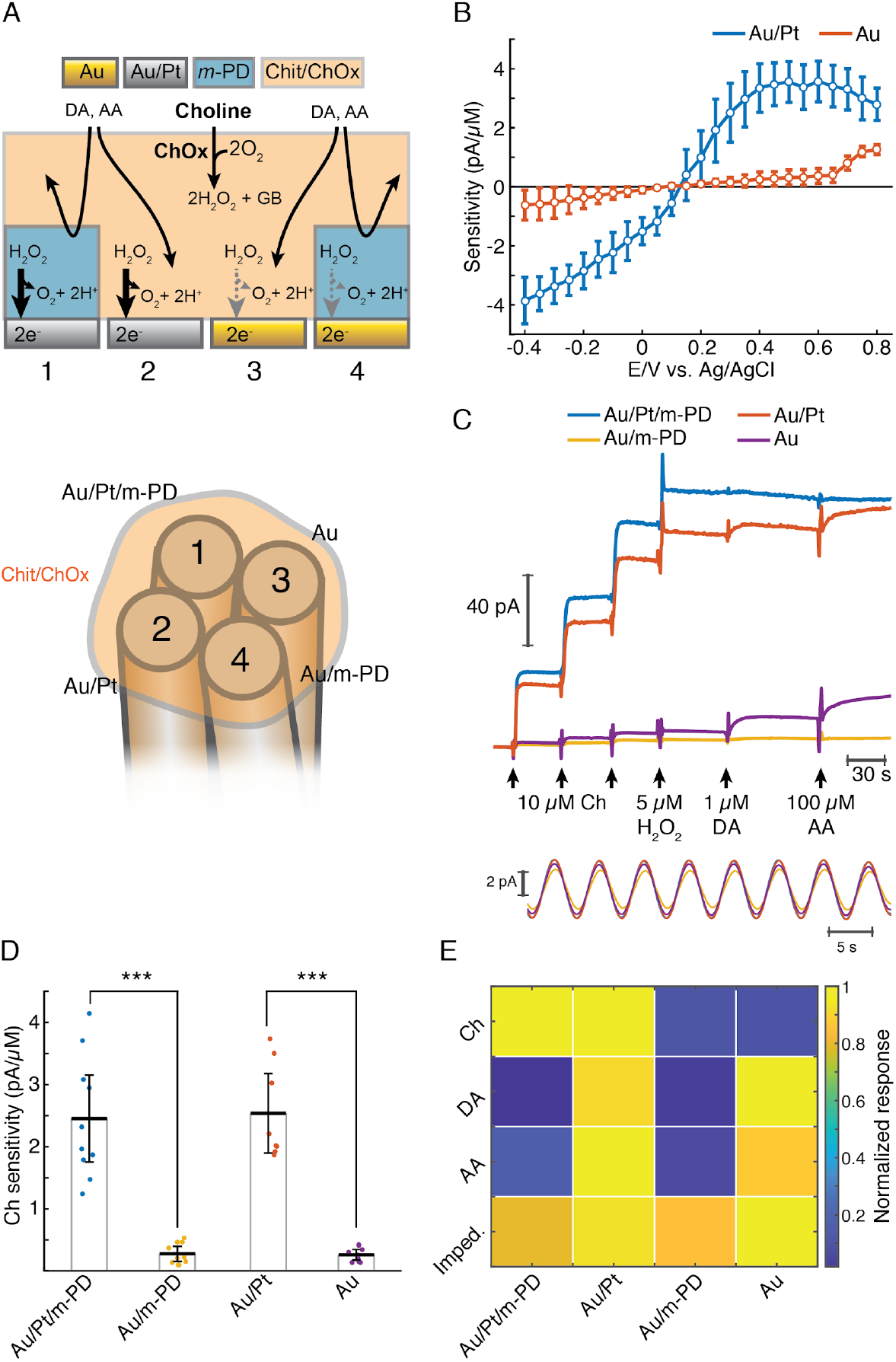
TACO sensor response properties and application in behaving animals. (A) Schematics depicting the multichannel biosensor design. (B) Voltammogram showing H_2_O_2_ sensitivities of gold-plated and platinized sites upon amperometric calibrations at different DC potentials (n=10). Prior to calibrations, the tetrodes were coated with a matrix of chitosan/ChOx. (C) Top shows a representative calibration of a sensor showing the response of different types of sites to step additions of Ch, H_2_O_2_, dopamine (DA) and ascorbate (AA). Bottom shows an example of current responses of sensor sites to a sinusoidal 12 mV AC voltage at 0.2 Hz overlaid on top of +0.6 V vs. Ag/AgCl DC voltage. The current AC amplitudes were used to calculate electrodes’ impedances at 0.2 Hz. (D) Sensitivities of different types of sites towards Ch (n=10 biosensors). Unlike *m*-PD electropolymerization, platinization significantly increased sensitivity (p<0.0001 and *F*=115 for platinization effect and p=0.87 and *F*=0.03 for *m-*PD, by two-way ANOVA for unbalanced data). (E) Normalized responses of each tetrode site to Ch, interferent molecules and voltage, presented as impedance at 0.2 Hz (n=5-10). Magnitudes significantly depended on the site modification and on the factor tested (p<0.0001 and *F*=22 for platinization, *F*=109 for *m*-PD and *F*=12.9 for factors, by three-way ANOVA for unbalanced data). Platinization selectively increased responses to Ch (p<0.0001) while *m*-PD decreased responses only to DA and AA (p<0.0001). Impedances did not significantly differ across different types of electrode modifications (p>0.99). Groups were compared by three-way ANOVA followed by Tukey-Kramer post-hoc tests.

The responses of TACO sensors’ sites to Ch, H_2_O_2_ and to compounds that can potentially interfere during *in vivo* measurements were tested by step additions in the beaker at +0.6V vs. Ag/AgCl (Figure 1C, top). In accordance with the voltammograms of H_2_O_2_ sensitivities (Figure 1B), the responses of TACO sensors’ gold-plated sites to Ch and H_2_O_2_ were much lower than those of platinized sites. On average, like for H_2_O_2_, this difference was about 10-fold for Ch, regardless of *m*-PD electropolymerization (Figure 1D). Contrasting with its lack of effect on Ch sensitivity, *m*-PD dramatically decreased responses to ascorbate and dopamine, regardless of the site’s metal composition (Figure 1E). In addition, we have also calibrated the impedance of recording sites at low frequency by applying a low-amplitude 0.2 Hz AC voltage on top of the DC offset. This low frequency is of particular relevance, as it overlaps with putative phasic cholinergic dynamics previously reported by us and other groups in anesthetized and freely-moving rodents (Howe et al., 2017; Parikh et al., 2007; Santos et al., 2015; Teles-Grilo Ruivo et al., 2017). Impedances calculated from the current oscillations generated by the AC voltage (Figure 1C, bottom) were comparable across all sites (Figure 1E). Collectively, the results summarized in Figure 1E and Table 1 validate the gold-plating approach to produce *pseudo*-sentinel sites, as it selectively reduces electrode’s response to H_2_O_2_. As compared to our previous stereotrode design using 50 μm diameter wires, these sensors keep the same *in vitro* performance, with a limit of detection (LOD) in the low nanomolar range and *T_50_* response times around 1.5 s (Table 1), which is remarkable for such small electrode surfaces. Additionally, while *m*-PD abrogates the response of electropolymerized sites to large electroactive molecules, the differential site modifications might provide further information on the signal identity. Allied to multivariate methods of analysis, the differential sites’ responses to different factors can potentially be exploited and electrochemical tetrode designed employed in TACO sensor can be generalized to other types of sensors.

**Table 1.** Analytical properties of TACO sensors

| Individual sites' analytical properties |  |  |  |  |  |  |
| --- | --- | --- | --- | --- | --- | --- |
| Channel type | Ch sensitivity<br>(pA/μM) | Ch sensitivity<br>(nA μM<sup>-1</sup> cm<sup>-2</sup>) | H<sub>2</sub>O<sub>2</sub><br>sensitivity<br>(pA/μM) | DA sensitivity<br>(pA/μM),<br>selectivity<br>ratio | AA sensitivity<br>(pA/μM),<br>selectivity<br>ratio | Impedance<br>(GΩ) |
| Au/Pt-mPD | 2.45 ± 0.70<br>(n = 10) | 1081 ± 309<br>(n = 10) | 2.48 ± 0.93<br>(n = 8) | 0.15 ± 0.23,<br>> 4.6 (n = 10) | 0.02 ± 0.013,<br>> 53 (n = 10) | 2.90 ± 0.48<br>(n = 5) |
| Au/Pt | 2.54 ± 0.64<br>(n = 8) | 1118 ± 283<br>(n = 8) | 2.86 ± 1.10<br>(n = 6) | 7.48 ± 1.86,<br>> 0.2 (n = 8) | 0.22 ± 0.20,<br>> 4.4 (n = 8) | 3.38 ± 0.56<br>(n = 5) |
| Au-mPD | 0.27 ± 0.12<br>(n = 10) | 118.2 ± 53<br>(n = 10) | 0.22 ± 0.16<br>(n = 8) | 0.13 ± 0.23,<br>> 0.41 (n = 10) | 0.007 ± 0.004,<br>> 14 (n = 10) | 3.11 ± 0.77<br>(n = 5) |
| Au | 0.25 ± 0.086<br>(n = 9) | 111.5 ± 38<br>(n = 10) | 0.29 ± 0.08<br>(n = 7) | 8.24 ± 1.84,<br>> 0.017 (n = 9) | 0.19 ± 0.16,<br>> 0.48 (n = 9) | 3.65 ± 0.59<br>(n = 5) |
| Whole sensor's analytical properties |  |  |  |  |  |  |
| Limit of detection (nM) |  |  | Response time (s) |  |  |  |
| 28 ± 0.011 (n = 10) | | | 1.4 ± 0.4 (n=10) | | | |

The data are given as the mean ± CI (95%)
The number of sensors tested is given in parentheses.

### Application of the TACO sensor in freely-moving rodents highlights the benefits of optimal suppression of confounding sources

In order to test the capacity of the TACO sensor to achieve artifact-free measurements of fast COA transients in behaving animals, we have first performed recordings in freely-moving animals. We implanted a TACO sensor in the rat brain, which was progressively lowered until it reached the hippocampal CA1 pyramidal layer, using a microdrive. Since the measurement of fast and local LFP-related patterns could potentially be affected by the enzyme coating, we concurrently recorded LFP from a 32-channel linear silicon probe implanted in the proximity of the biosensor. Extracellular electrophysiology allowed us to validate the amperometric measurement of high-frequency LFP-related profiles while gathering multi-layer hippocampal network dynamics (Figure 2A, left).

**Figure 2.**
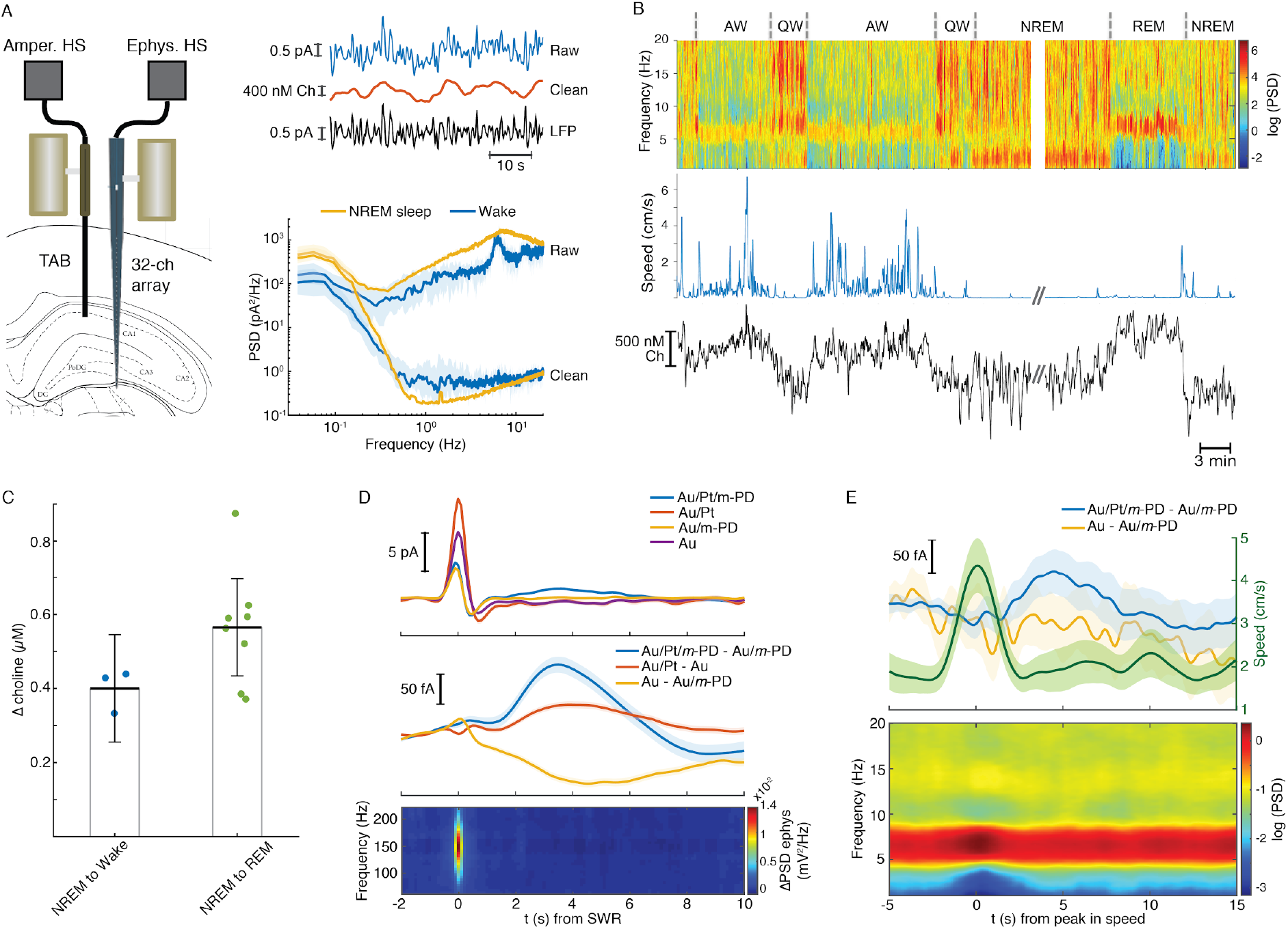
TACO sensor application in freely-behaving animals. (A) Left panel depicts the arrangement of a multichannel ChOx biosensor and a 32-channel silicon probe chronically implanted in the hippocampus of a rat. Both probes were attached to microdrives. Top right panel shows a segment of a raw signal (low-pass filtered at 1 Hz) and the resulting clean putative Ch and LFP components upon common-mode rejection, recorded during NREM sleep. The plot in the bottom right illustrates the dramatic decrease in the spectral power upon cleaning the signal by common-mode rejection both during wake (n=10) and NREM sleep periods (n=19). (B) Representative recording across multiple brain states showing power spectrogram from a silicon probe site at hippocampal fissure (top), rat speed (middle) and clean ChOx signal (bottom). (C) Change in tonic putative Ch levels following transitions from NREM sleep to wake (n=3) or REM sleep (n=9). Both groups were significantly different from zero (p<0.05, inferred from one-way anova followed by Tukey-Kramer post-hoc analysis). (D) Average low frequency (1 Hz low-pass filtered) biosensor signals and high frequency power spectrograms triggered to SWRs detected from a silicon probe channel in CA1 pyramidal layer (n=8074). Responses from biosensor sites are shown before (top) and after (middle) common-mode rejection. Average spectrogram was obtained from the silicon probe site used to detect SWRs (bottom). (E) Average speed, biosensor signals following common-mode rejection and LFP power spectrogram triggered to peaks in rat speed in an open-field arena (n=127, top). The spectrogram was derived from hippocampal fissure LFP (bottom).

The COA signal was cleaned by subtraction of the *pseudo*-sentinel from the Ch-sensing sites’ signal, upon frequency-domain correction of the *pseudo*-sentinel amplitude and phase (please see Methods) (Santos et al., 2015). This procedure led to substantial removal of fast current fluctuations ascribed to LFP (example recording in Figure 2A, top right). Accordingly, the mean spectral power at ~1-20 Hz of the signal derived from the Au/Pt/m-PD site during both NREM sleep and wake periods decreased more than two orders of magnitude after the signal cleaning procedure (Figure 2A, bottom right). The cleaned COA signal tonically changed across different brain states reaching higher values during active wakefulness and REM sleep than during NREM sleep, which corroborates previous measurements of brain state-dependent ACh levels (Hasselmo and McGaughy; Marrosu et al., 1995) (Figure 2B and C). Remarkably, on top of tonic brain-state-related dynamics, COA fluctuated on the time-scale of seconds, suggesting the possibility of a rich phasic cholinergic dynamics within each brain state.

Acetylcholine can suppress the occurrence of SWRs (Norimoto et al., 2012; Vandecasteele et al., 2014) and positively correlates with arousal (Hasselmo and McGaughy; Marrosu et al., 1995; Reimer et al., 2016; Teles-Grilo Ruivo and Mellor, 2013; Teles-Grilo Ruivo et al., 2017), which typically coincides with hippocampal theta oscillations and manifests as locomotion in freely-moving animals (Buzsáki, 2002; Gu et al., 2017). Thus we tested whether these events could correlate with fast changes in putative cholinergic activity reflected by COA. hippocampal SWRs were detected from a silicon probe site in the CA1 pyramidal layer during NREM sleep. Average raw biosensor signals triggered on the peak of SWRs showed a prominent peak in all TACO sensor’s sites (Figure 2D, top). The similarity of peak amplitudes suggests an LFP-related origin of these currents, which was virtually absent in the cleaned signals (Figure, 2D, middle), and reflects the slow profile of the sharp wave. The high magnitude of this LFP signal contaminating COA measurements emphasizes the importance of the common-mode rejection approach in revealing putative cholinergic dynamics that would otherwise be masked. Interestingly, regardless of *m*-PD electropolymerization, putative Ch differential signals showed a peak lagging the SWR by ~3 s. Importantly, this peak was not present in *pseudo*-sentinel sites, supporting an authentic change in COA (Figure 2D, top and middle).

In our recordings, high-frequency bursts captured with electrophysiology and amperometry typically coincided in time and exhibited similar spectral features (Figure S2A). Accordingly, the continuous power-power correlation between amperometry-derived current and silicon probe-derived LFP was high (>0.7, Figure S2B) across a wide frequency range. The very similar average power spectra triggered to SWRs (Figure 2D, bottom and Figure S2C) and the high co-occurrence of SWRs detected with both modalities (Figure S2D) further prove the ability of our amperometric system to reliably record high-frequency LFP-related currents. Although theoretically expected, this capability has not been experimentally documented before.

Bouts in locomotion, detected as peaks in the rat running speed, were associated with transient increases in theta power and were also followed by a peak in putative Ch signal, observed in the clean biosensor signal but not in the *pseudo*-sentinel subtraction control (Figure 2E).

These results highlight the usefulness of our multichannel *pseudo*-sentinel approach to discriminate between authentic changes in COA and interferent signals in the brain. Notably, phasic changes in COA were associated with transient changes in arousal or exploratory behavior and with SWRs, events that are critical for memory encoding and consolidation, respectively (Buzsáki, 2002, 2015).

### Phasic biosensor responses during locomotion bouts correlate with oxygen transients

TACO sensor offers the opportunity to study cholinergic activity with unprecedented spatial resolution and selectivity in behaving animals. However, since O_2_ is a co-substrate of oxidases, sensitivity to physiological O_2_ variations is a potential issue of this type of biosensors (Baker et al., 2015; Chatard et al., 2018; Dixon et al., 2002; McMahon et al., 2007; Santos et al., 2015). To control for O_2_-related signals *in vivo,* we have exploited the advantages of the tetrode configuration, ideally suited to measure multiple electroactive compounds at the same brain spot. Furthermore, bias on the impedance and response time across the recording sites is minimal since the same enzyme immobilization matrix covers the whole tetrode tip. Using a new customized head-stage allowing independent control of the potential applied on each recording site, we have simultaneously measured O_2_ and COA with the same sensor. Oxygen measurement was achieved by O_2_ reduction on a gold-plated site at a negative potential, which was typically maximal at −0.4 V vs. Ag/AgCl (Figure 3A, left). To avoid electrical cross-talk between tetrode sites, sporadically observed when we applied a high negative voltage to one site, we set −0.2 V vs. Ag/AgCl as the standard voltage for *in vivo* O_2_ measurements. Although the gold-plated site could also reduce H_2_O_2_, the very low response magnitude as compared to O_2_ (Figure 3A, left) assured a negligible contribution of H_2_O_2_ generated by immobilized ChOx to the O_2_ signal recorded *in vivo*. Under this configuration (Figure 3A, bottom), clean COA and O_2_ signals were obtained by subtraction of Au/Pt//*m*-PD (at +0.6 V) and Au (at −0.2 V) by the *pseudo*-sentinel /*m*-PD site, respectively.

**Figure 3.**
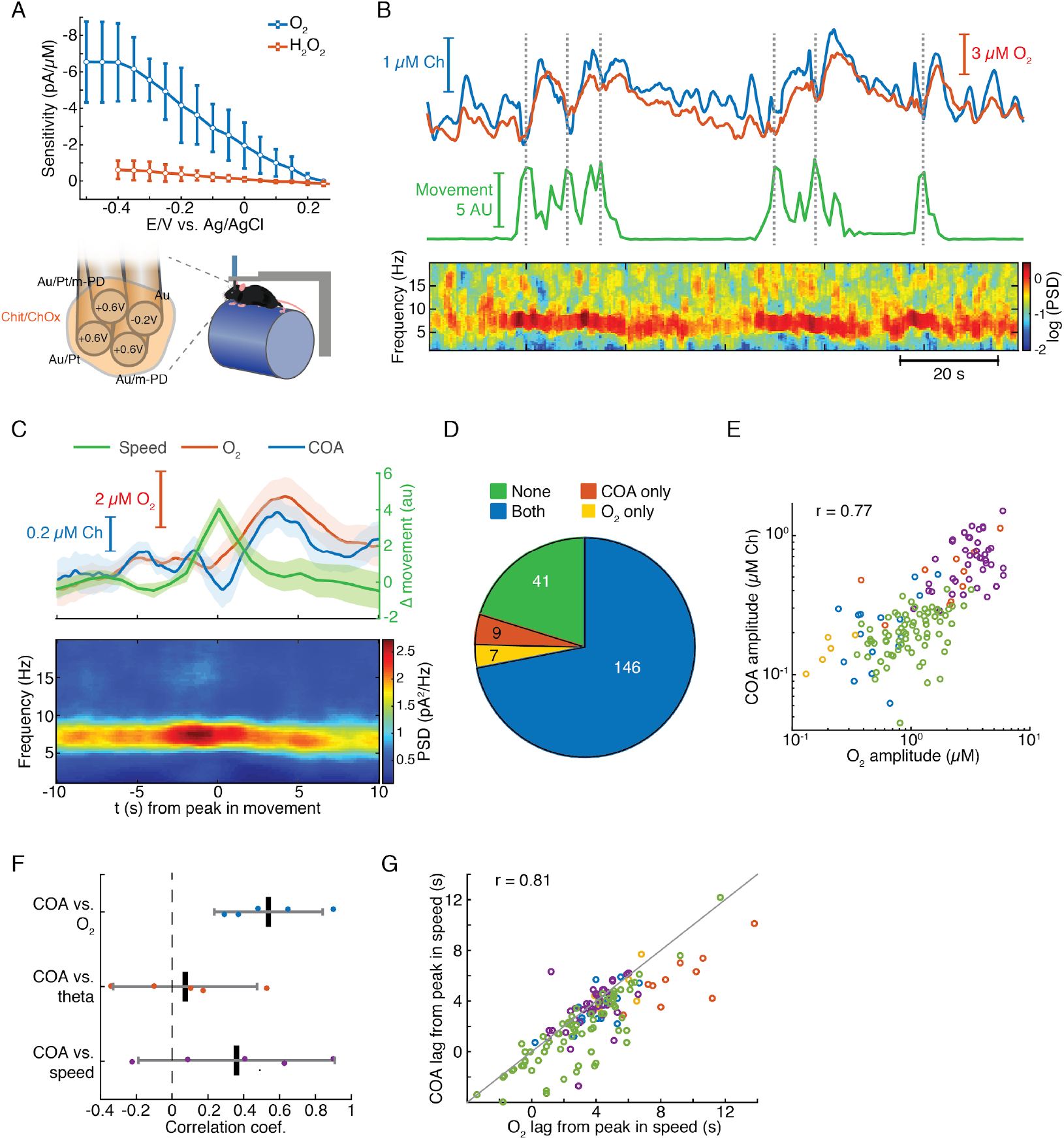
Locomotion-related correlated changes in COA and local oxygen concentration in head-fixed mice. (A) The voltammogram shows the DC voltage-related sensitivity of gold-plated sites towards H_2_O_2_ and O_2_ (top). The schematics shows the tetrode configuration used to simultaneously measure H_2_O_2_ resulting from COA and extracellular O_2_ (bottom left). Values on each recording site indicate the applied DC voltage *vs.* Ag/AgCl (+0.6 V for H_2_O_2_ and −0.2 V for O_2_ measurements). Schematics of the head-fixed setup (bottom right). (B) Representative data segment of simultaneous recording of COA and O_2_ (top), locomotion speed (middle) and LFP spectrogram(bottom). Dashed lines indicate the times of detected locomotion bouts. (C) Average speed, COA and O_2_ profiles (top) and LFP power spectrogram bottom) triggered to locomotion bouts (n=41 from one recording session). (D) Event counts across different categories, distinguished by the occurrence or absence of ChOx and/or O_2_ peaks following locomotion bouts. Data was collected from 5 recording sessions in 3 mice. (E) Amplitude of ChOx (shown as calibrated Ch concentrations) vs O_2_ transients following locomotion bouts (data from the events that had increases in both signals, n=146). Each color represents one recording session (n=5-74 events per recording). Amplitudes were significantly correlated (r_spearman_ = 0.77, p<0.001). (F) Spearman correlation coefficients between ChOx activity amplitudes and O_2_, theta power or speed (n=5). Correlation across recordings was significantly above zero for ChOx *vs* O_2_ data (p<0.01, *t*-test). Each point represents a single recording session. (G) Lags of ChOx peaks *vs.* O_2_ peak lags relative to the peak in speed. The correlation between variables was significant (r_spearman_=0.81, p<0.001, n=146). Each color represents one recording session (n=5-74 events per recording). The diagonal line was plotted to ease the comparison between lags.

In order to obtain more controlled experimental conditions and overcome technical constraints posed by the size of the head-stage, the remaining experiments were performed in head-fixed mice spontaneously running on a treadmill (Figure 3A bottom). Strikingly, we found that phasic COA dynamics typically matched the simultaneously recorded O_2_ fluctuations, which were generally related to changes in behavioral state (Figure 3B). In line with the freely-moving data (Figure 2E), head-fixed running bouts were temporally correlated with an increase in the power of theta oscillations as well as with delayed phasic increases in both COA and O_2_, peaking a few seconds later (Figure 3B and C). This lag was significantly greater than zero in all recording sessions (two-sample *t*-test, p<0.01), averaging 3.85 ± 2.04 s (n = 5) for COA and 5.04 ± 3.12 s (n = 5) for O_2_. Notably, running-related isolated COA or O_2_ peaks were rare, with the vast majority of the events showing either no identifiable change or co-occurrence of the two transients (Figure 3D). Amplitudes of COA peaks were significantly correlated with those of O_2_ (p<0.001) when pooling together the events from all recordings, which allowed sampling across the full O_2_ amplitude range (Figure 3E). Phasic COA correlated more consistently with O_2_ than with theta power or speed (Figure 3F). Moreover, runningbout-related COA and O_2_ peak lags were significantly correlated (p<0.001, Figure 3G) and, interestingly, within most sessions (3 out of 5), COA peaked significantly earlier than O_2_ (p<0.05). In summary, these results indicate a strong correlation between phasic COA and O_2_ in the hippocampus of head-fixed mice following locomotion bouts.

### Phasic COA and oxygen signals follow clusters of sharp-wave/ripples during immobility

Hippocampal SWRs are critical for memory consolidation and their occurrence has been proposed to be anti-correlated with cholinergic activity in the hippocampus (Hasselmo and McGaughy; Norimoto et al., 2012; Vandecasteele et al., 2014). However, our freely-moving data showing COA transients following SWRs contradicts this prediction, posing questions on the factors driving the biosensor response during these events. Thus, we investigated whether the SWR-related response of immobilized ChOx was correlated with extracellular O_2_ in head-fixed mice during periods of quiescence.

Remarkably, on average, SWR events were followed by fast transients in both COA and O_2_ (Figure 4A). Both COA and O_2_ peak amplitudes correlated best with ripple power integrated over a period of ~2 seconds lagging them by 3-4 seconds (Figure 4B and C). Similarly, both SWR count and summed ripple power integrated in a 2-second window positively correlated with the delayed amplitude of both COA and O_2_ transients (Figure 4D-E and Figure S3).

**Figure 4.**
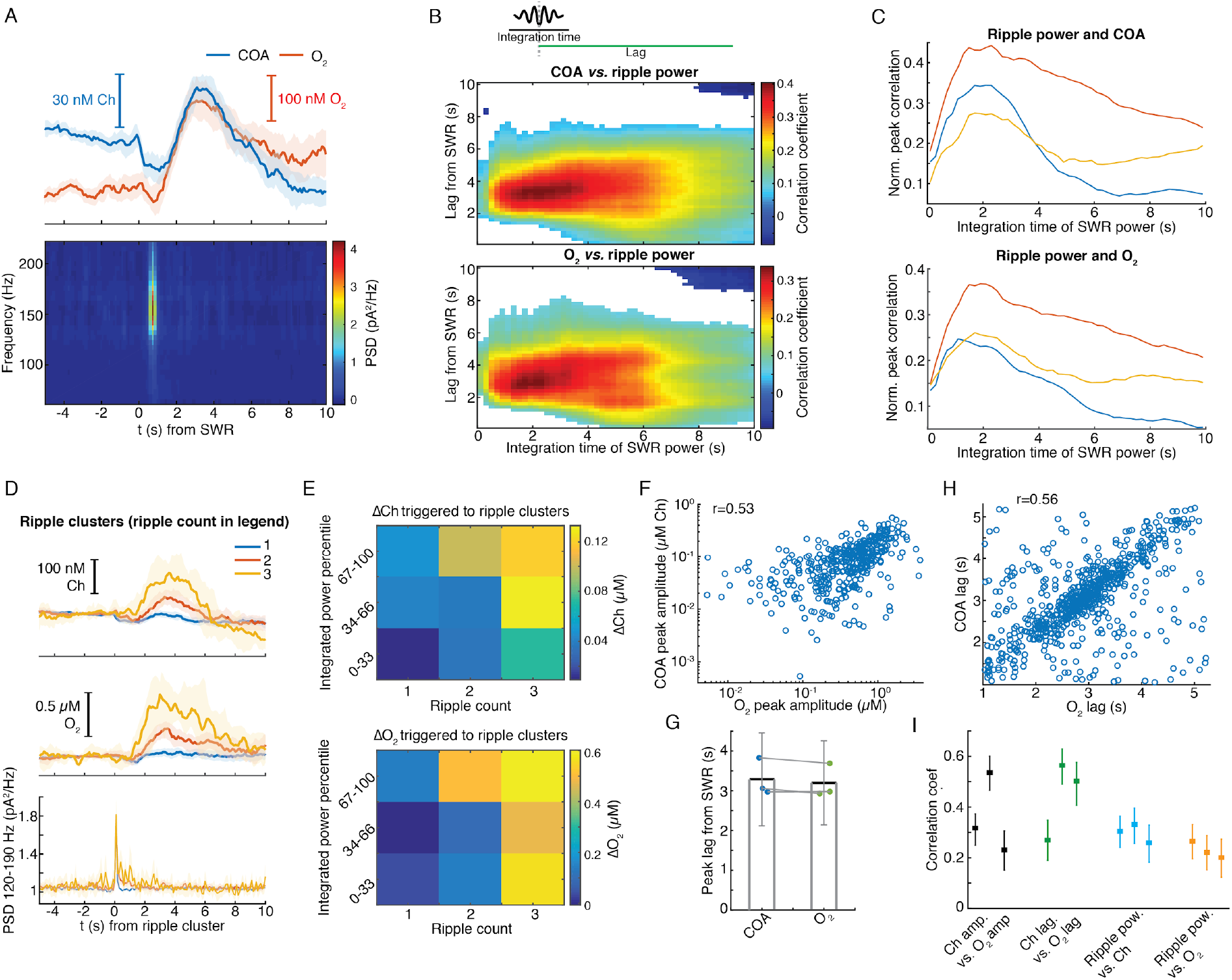
Correlated COA and oxygen dynamics associated with hippocampal SWRs. (A) Average ChOx and O_2_ dynamics triggered to hippocampal SWRs (n=1067) in the head-fixed setup (top) and average LFP spectrogram (bottom). (B) Pseudo-color-coded Spearman correlation between ChOx (top) or O_2_ (bottom) amplitude at different time lags from SWRs (y-axis) and integrated ripple band power computed in windows of varying size (x-axis) for a representative session. White areas represent non-significant correlations (p>0.05). (C) Normalized peak correlations, obtained from the difference between maximal and minimal correlations for each integration time in B, between integrated ripple power and ChOx activity or O_2_. Colors represent recording sessions in different mice. (D) Average COA, O_2_ and ripple power dynamics triggered to SWRs bursts containing variable number of SWRs (1-3) in a 2 s window (n=34-591 SWRs in each group, respectively from one representative session). (E) Peak amplitude of ChOx activity or O_2_ transients as a function of SWR burst size (as shown in D) sorted by different percentile ranges of summed ripple power (same data as in D). Both SWR count and total ripple power significantly affected the amplitude of ChOx and O_2_ (two-way ANOVA for unbalanced data following ART, *p*<0.0001 and *F*>12 for both factors in ChOx and O_2_ data). (F) Amplitude of ChOx *vs.* O_2_ transients following SWRs from one recording session (r_spearman_= 0.53, p<0.0001, n=1067). (G) Group statistics on the lags of ChOx and O_2_ peaks relative to SWRs. Each dot is the average from one recording. Bars represent means ± CI. (H) Lags of ChOx peaks as a function of O_2_ peak lags relative to SWRs. The parameters were significantly correlated (r_spearman_=0.56, p<0.0001, n=1067). (I) Summary of correlations between sensor signals and ripples. Each point represents one recording session, and error bars are CIs computed using bootstrap.

These findings might indicate the contribution of a time-constant related to the sensor response and/or to a relatively slow physiological process by which SWRs recruit cholinergic activity or a local hemodynamic response leading to O_2_ increase. Indeed, functional MRI has reported SWR-triggered increases in BOLD signal in the primate hippocampus, reflecting a local tissue hemodynamic response at a time-scale matching O_2_ transients observed here (Ramirez-Villegas et al., 2015). Importantly, the amplitudes of SWR-associated COA and O_2_ phasic transients were consistently correlated within all recordings (n=3, p<0.001, Figure 4F). Similarly, lags of these transients to SWR were significantly correlated (Figure 4G, H, I).

Together, the data indicate correlated phasic profiles of COA and extracellular O_2_ in response to SWRs (Figure 4I), especially when they happen in clusters. As in the case of locomotion bouts, at this stage one could not rule out the contribution of neither phasic Ch or O_2_ as the trigger for the peaks in COA. The putative cholinergic origin of this response would, nevertheless, be surprising given the suppressive effect of ACh on SWR occurrence (Norimoto et al., 2012; Vandecasteele et al., 2014). Thus, in light of the O_2_ transients that accompany the rise in COA, these observations cast doubt on the validity of the putative SWR-triggered cholinergic response.

### Interactions between COA and oxygen are not sensitive to ongoing hippocampal dynamics and depend on the time-scale

The correlation between COA and O_2_ following locomotion and SWRs may reflect interaction between the two signals or result from the coincident recruitment of cholinergic and hemodynamic responses. While a consistent relationship between the two variables is expected in the first case, irrespective of ongoing network dynamics, the same may not happen in the latter. To get insights into this question, we analyzed an additional category of events, consisting of fast O_2_ transients detected outside the time-windows surrounding SWRs and peaks in locomotion. Under this condition, the average rate of O_2_ peaks occurrence was only 18% of the rate computed from total O_2_ peaks (Figure S4A), emphasizing the strong modulatory effect of hemodynamics and respiration on O_2_, potentially evoked by SWRs and locomotion, respectively (Leithner and Royl, 2014; Ramirez-Villegas et al., 2015; Zhang et al., 2019). Remarkably, virtually all (>95%) of these events had an associated COA transient. The dynamics of COA and O_2_ were typically similar (Figure 5A), with COA peaks lagging, on average, from −0.09 s to 0.48 s (n=3 recordings) relative to O_2_. Signal amplitudes were significantly correlated both when pooling together all events (p<0.001, Figure 5B) or within all individual sessions (p<0.01). The amplitude correlation coefficients were in the range of those obtained for O_2_ peaks associated with locomotion and SWRs (Figure 5C), but the average amplitudes of O_2_ and corresponding COA signals were in the sub-micromolar range, comparable to those associated with SWRs (Figure 5D). Overall, although the amplitude range of locomotion-related peaks was wider, it is notable that the whole data fit to a COA/O_2_ relationship that seems to follow the same model across recordings and event types (Figure 5D). These observations are therefore compatible with an interaction between COA and O_2_, although contribution of third-party factors could not be definitely excluded at this stage.

**Figure 5.**
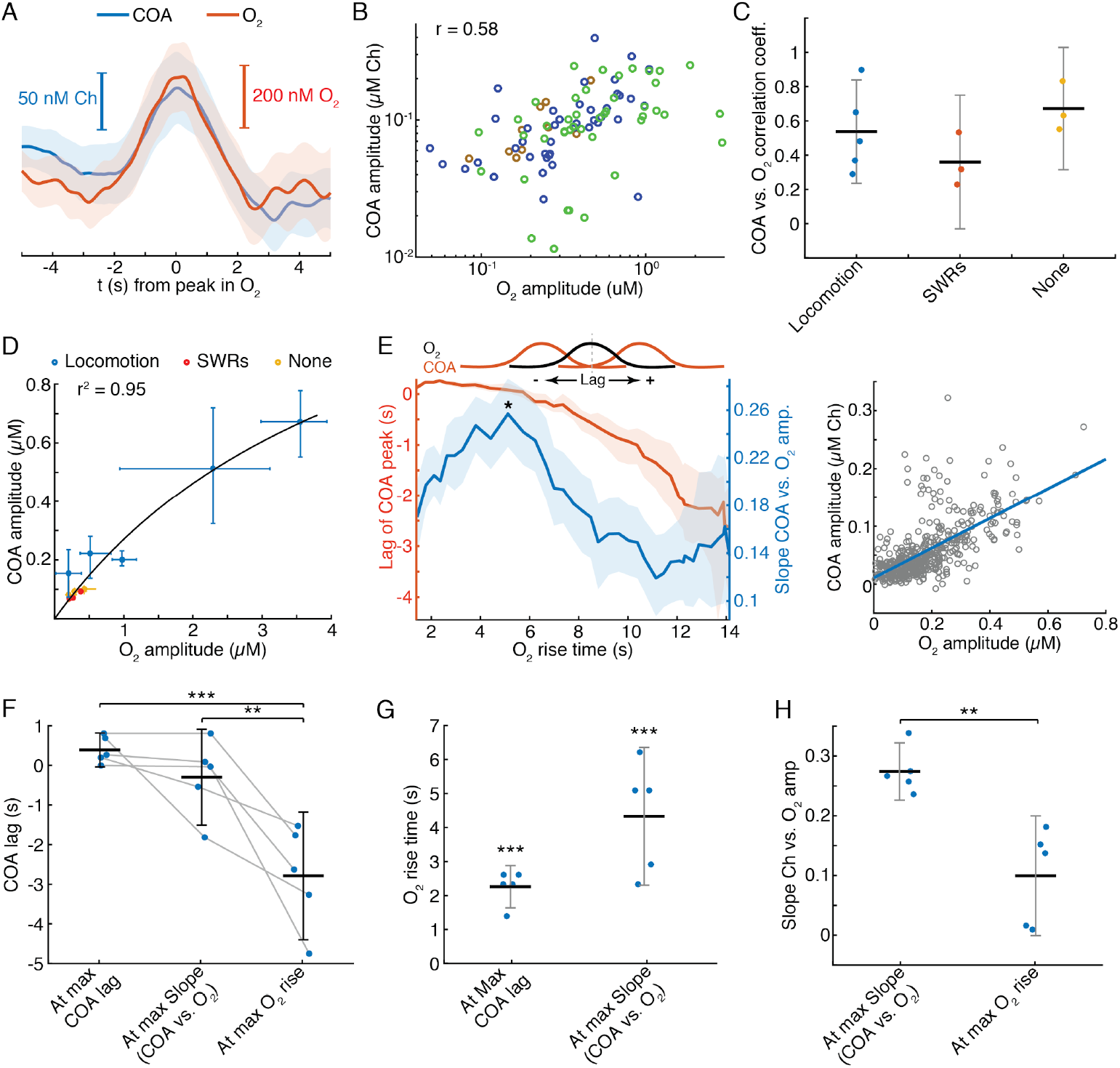
Correlation between COA and spontaneous O_2_ transients. (A) Average COA and O_2_ dynamics triggered to fast O_2_ transients (duration ~5 s) detected outside periods when SWRs or locomotion bouts occurred. Data is from one recording session (n=42 events). (B) Relationship between amplitudes of O_2_ and associated COA transients outside SWRs/locomotion bouts. Colors represent different recording sessions (n=10-45 from 3 recordings, r_spearman_=0.58, p<0.001). (C) Group summary of COA *vs.* O_2_ amplitude correlations under different behavioral and/or electrophysiological contexts. (D) COA vs. O_2_ amplitude across animals and states. Each point represents the median of events from a recording session. Data were fitted to the Michaelis-Menten equation, resulting in a Vmax = 1.58 and a Km = 4.84. (E) Lag of COA peaks relative to O_2_ peaks (red) and slope of COA *vs.* O_2_ amplitude (blue) for O_2_ transients with varying rise time for a single session; medians with CIs (left). Right shows the relationship between amplitudes of O_2_ and associated COA transients for the O_2_ rise time corresponding to largest slope marked with * on the left panel. (F) Group statistics on maximal COA lags, lags at maximal COA/O_2_ slope and lags associated with longest O_2_ transients (ANOVA, F = 15.22, with post-hoc Tukey test, p=0.51 for max COA lag *vs*. lag at max COA/O_2_). (G) Group statistics on O_2_ rise times corresponding to maximal COA lags and COA/O_2_ slope in (E). Values from both groups were significantly lower than the longest O_2_ rise time observed in a given recording (p<0.001, one-sample *t*-test). (H) Group statistics on the slopes of COA vs. O_2_ amplitude at its maximum value and at maximum O_2_ rise. Differences between groups were significant (p<0.005, paired *t*-test). *** p<0.001, ** p<0.01.

Besides the amplitude of O_2_ transients, the temporal profile of O_2_ rise might influence the shape and amplitude of associated ChOx responses and thus provide further hints on the causality and directionality of COA-O_2_ interactions. Thus, we detected O_2_ peaks at different frequency bands (ignoring their correlation with SWRs or speed), resulting in a spectrum of O_2_ rise times from less than 2 to 14 s (please see *Methods* section for details). Across all recording sessions, we consistently observed that the COA peak anticipated O_2_ (negative lag) as O_2_ rise time increased (Figure 5E,F and Figure S4B). Maximal COA lags averaged 0.38 ± 0.42 s and were associated with fast O_2_ rises, lasting 1.4-2.6 s (Figure 5F and G). Furthermore, the slope of COA vs. O_2_ amplitudes was time-scale dependent, peaking for O_2_ transients that took 2.3 to 6.2 s to rise (Figure 5E, G and Figure S4C) and progressively decreasing for longer O_2_ rises (Figure 5E, H and Figure S4C). The time-scale dependence of the COA vs. O_2_ slope is apparently related to the non-linear interaction between COA and O_2_, as a function of O_2_ rise time. Contrasting with the approximately linear increase in O_2_ amplitude as a function of O_2_ rise, the amplitude of corresponding COA peaks was, on average, nearly time-scale independent for O_2_ rise times in the range of 3-14 s (Figure S4D).

In general, these results provide important insights into the interaction between COA and extracellular O_2_ in the hippocampus. The advancement of COA relative to O_2_ is apparently compatible with a Ch-O_2_ directionality, possibly caused by ACh-evoked changes in local blood flow (Takata et al., 2013). However, the diffusional delay underlying such a mechanism is expected to be constant as a function of time-scale. Moreover, the time-scale dependence of the relative amplitude of enzyme response is hard to interpret in light of a Ch-O_2_ directionality. Alternatively, non-steady state ChOx responses to O_2_ transients are fully compatible with our observations. Such modulation of sensor response is, by definition, phasic and is expected to be amplified within an optimal, probably short, time-window. Importantly, since phasic responses are short-lived, in addition to the time-scale dependence of magnitude, this phenomenon predicts a continuous drift in COA-O_2_ peak lags as O_2_ rise time increases.

Together, these data converges to the hypothesis that the observed changes in COA are caused by O_2_ fluctuations.

### Exogenous oxygen transients in the hippocampus elicit phasic COA responses

Our correlational analysis points towards a possible effect of O_2_ transients on phasic sensor responses in the hippocampus. We tested this hypothesis first by evoking changes in O_2_ by local application of small volumes of O_2_-saturated saline through a glass micropipette, positioned at a few hundred microns from the biosensor tip. To evoke different O_2_ profiles, we varied ejection parameters such as time and pressure.

Remarkably, immobilized ChOx exhibited robust phasic responses to exogenous O_2_ transients, regardless of the time-scale of O_2_ change (Figure 6A and Figure S5A). Amplitudes of COA and O_2_ were significantly correlated (p<0.0001, Figure 6B). However, similarly to spontaneous dynamics, the CAO/O_2_ amplitude ratio of single events (equivalent to CAO/O_2_ slope in spontaneous data) significantly decreased with O_2_ rise time in all experiments (negative correlation, p<0.05, Figure 6C). The decrease in COA amplitude was accompanied by the advancement of its peak relative to O_2_ (p<0.001 in all recordings, Figure 6D). Thus these results qualitatively recapitulate our observations for the spontaneous COA-O_2_ interactions, reinforcing the putative O_2_-COA directionality.

**Figure 6.**
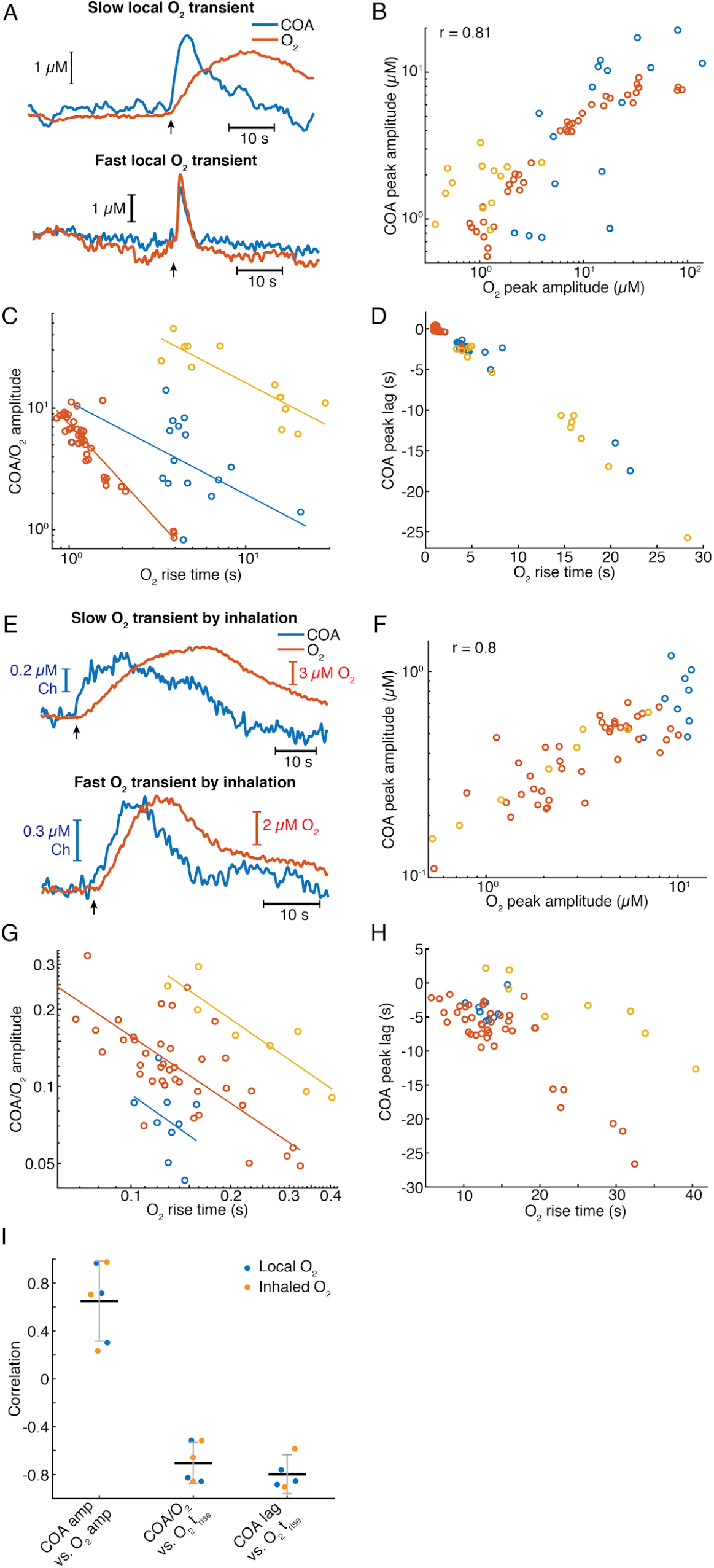
Exogeneous O_2_ elicits phasic COA responses in the hippocampus *in vivo.* (A) Representative examples of slow and fast O_2_ transients and associated COA responses evoked by local application of exogenous O_2_ from a glass micropipette. (B) Amplitudes of COA *vs.* locally-evoked O_2_ transients. Data were collected from three recording sessions (color-coded, n=13-40 per session). Amplitudes were significantly correlated when pooling all data (r_spearman_=0.81, p<0.0001) and within 2 sessions (p<0.005, in one session this analysis was not performed due to the narrow range of O_2_ amplitudes covered). (C) Ratio of COA *vs*. evoked O_2_ peak amplitudes as a function of O_2_ signal rise time. Colors denote recording sessions and trendlines are linear fits performed on data from each session. In all cases, the variables were negatively correlated (p<0.05). (D) Lag of COA relative to locally-evoked O_2_ transient peaks as a function of O_2_ rise time. Lags significantly decreased with O_2_ rise time in all sessions (r_spearman_ for each recording ranged from −0.76 to −0.88, p<0.001). (E) Representative traces showing slow and fast O_2_ transients and associated COA responses evoked by O_2_ inhalation. E-H plots are analogous to A-D, but for O_2_-inhalation-induced O_2_ transients. (F) Amplitudes of ChOx *vs.* O_2_ transients evoked by inhalation. Data were obtained from three recording sessions (color-coded, n=8-40 per session). Variables were significantly correlated both when pooling the whole data (r_spearman_= 0.8, p<0.0001) and within individual experiments (p<0.0005). (G) Amplitude ratio of COA *vs*. inhalation-evoked O_2_ transients as a function of O_2_ signal rise time. Colors denote recording sessions and trendlines are linear fits performed on data from each session. The variables were significantly negatively correlated (red and yellow sessions, with r_spearman_ coefficients of −0.66 and −0.86 respectively, p<0.05). (H) Lags of COA relative to inhalation-evoked O_2_ peaks as a function of O_2_ rise time. Lags significantly decreased with O_2_ rise time (red and yellow sessions, r_spearman_ of −0.58 and −0.90 respectively, p<0.005). The analysis in F-H was not performed within one session (blue dots) due to insufficient coverage of O_2_ amplitudes and rise times. (I) Summary of amplitude and COA-O_2_ lags correlations for COA and O_2_ transients evoked by local O_2_ application and O_2_ inhalation. Data are presented as means ± CIs.

Next, we manipulated hippocampal O_2_ levels non-invasively, through inhalation, to further confirm O_2_ causality. We exposed mice to a pure O_2_ stream during variable periods (4-30 s) in order to generate different O_2_ transients. Like in the case of local O_2_ delivery, we observed reproducible changes in COA in response to exogenous O_2_ (Figure 6E and Figure S5B). The data showed a significant correlation between peak amplitudes (p<0.0001, Figure 6F). Importantly, both the COA/O_2_ amplitude ratio and COA peak lag significantly decreased as a function of O_2_ rise time (p<0.05), corroborating the conclusions from local O_2_ manipulation.

Overall, O_2_ inhalation tended to generate slower and smaller O_2_ transients (and perhaps more physiological) than those by local application. Despite the differences in magnitudes and time-scales, the tested correlations are consistent across the two paradigms (Figure 6I) as well as with the spontaneous interactions between COA and O_2_ (Figure 5E-H). The results converge to the hypothesis that the putative cholinergic transients observed in the hippocampus of behaving rodents rather result from phasic modulation of COA by physiological O_2_ fluctuations.

### Biosensor responses to oxygen have tonic and phasic components

Given the *in vivo* causality between O_2_ and COA, we sought a detailed investigation of the biosensor O_2_-dependence *in vitro,* in order to get mechanistic insights into the relationship between these signals. The *in vitro* tests were based on step additions of known O_2_ concentrations in the presence of a background Ch concentration (5 μM) representative of average brain extracellular Ch tonic levels (Brehm et al., 1987; Garguilo and Michael, 1996; Parikh et al., 2004). Unlike previous studies, this allowed us to get a clear distinction between phasic vs. steady-state (tonic) sensor responses.

Notably, upon removal of O_2_ from solution and in the presence of background Ch, most biosensors responded to consecutive O_2_ steps with a fast transient (phasic component) before reaching a steadystate (Figure 7A). Both phasic and tonic components decreased with O_2_ baseline, but not equally. The phasic response was usually still prominent upon the exhaustion of the tonic component (Figure 7A). We quantified these differences by fitting the Michaelis-Menten equation to the tonic changes and the Hill equation to the cumulative phasic peaks, as the latter did not appear to follow a pure Michaelis-Menten profile (Figure 7B). The resulting phasic K_0.5_O_2_ values for phasic responses were, on average, one order of magnitude larger than tonic KmO_2_, reinforcing that the phasic component vanishes at much larger O_2_ baselines than tonic responses (p<0.0005, Figure 7C). Importantly, the phasic ChOx response was not matched by a fast O_2_ transient following each addition, as O_2_ raised considerably slower than the peak in COA (Figure 7D).

**Figure 7.**
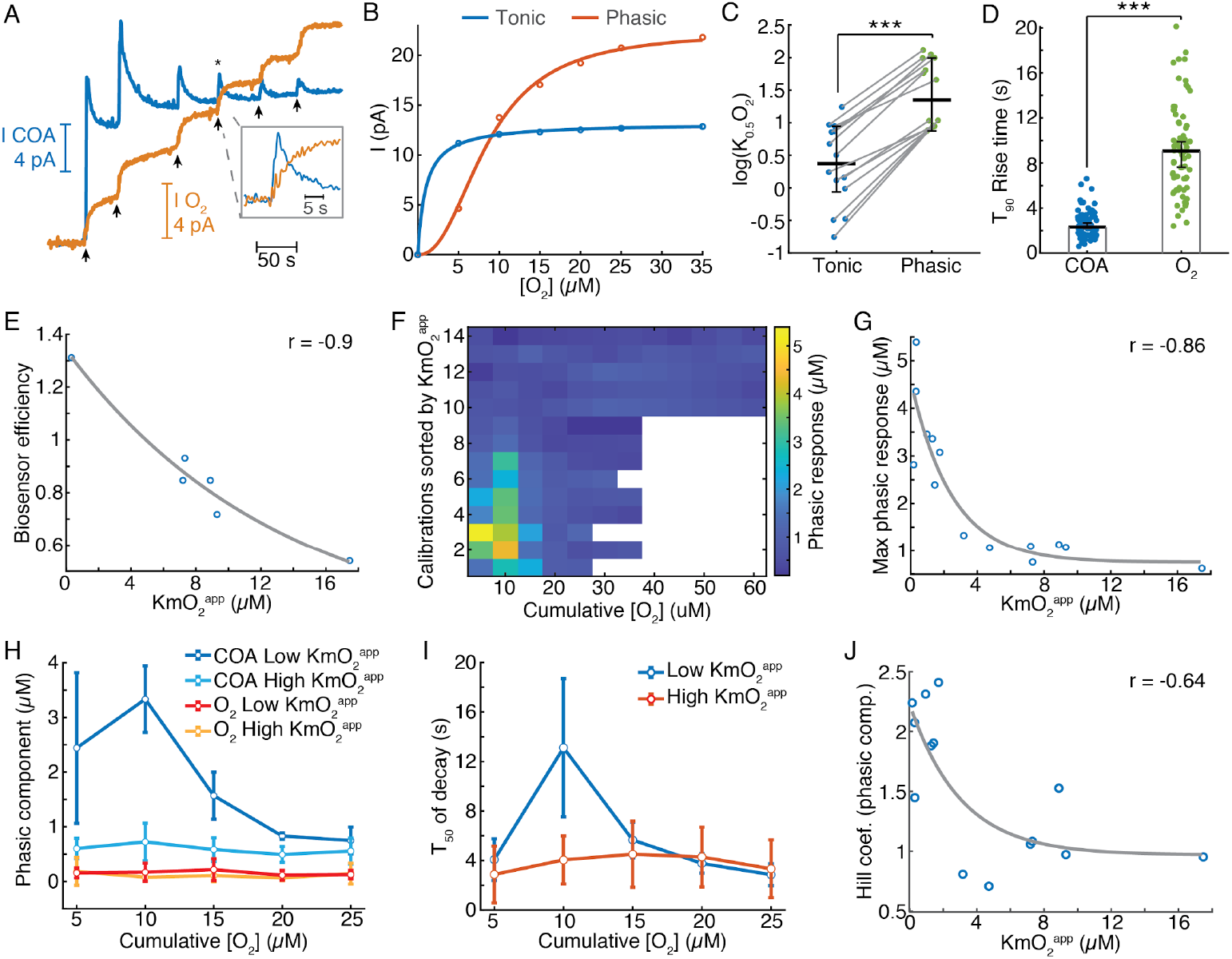
Biosensors generate tonic and phasic COA components in response to O_2_. (A) An example of an *in vitro* O_2_ calibration. Upon removal of O_2_ from PBS containing 5 μM Ch, step additions of 5 μM O_2_ (arrows) were performed until the tonic sensor response saturated. (B) Representative cumulative function of tonic and phasic responses as a function of O_2_ baseline after each addition. Tonic data were fitted with the Michaelis-Menten equation, resulting in a KmO_2_^app^ of 0.97 μM (CI = 0.818-1.126 μM) and I_max_ = 13.2 pA (CI = 13.04-13.37 pA), with RMSE = 0.08 pA. Phasic responses were fitted to the Hill equation, yielding K_0.5_O_2_ = 8.6 μM (CI = 7.58-9.62 μM), Hill coefficient *n* = 2.31 (CI = 1.74-2.88) and I_max_ = 22.4 pA (CI = 20.6 – 24.1), with rmse = 0.50 pA. (C) K_0.5_O_2_^app^ values of tonic and phasic components from all biosensors (n=14). Averages and error bars are medians and 95% CIs. The two groups were significantly different (sign test, p<0.0005). (D) T_90_ rise times of O_2_ steps and COA peaks following O_2_ additions (n=72-95). Bars and error bars represent medians and 95% CIs, two samples were significantly different (p<0.0001, Wilcoxon rank sum test). (E) Biosensor efficiency (Ch/H_2_O_2_ sensitivity ratio) as a function of KmO_2_^app^ (n=6). For illustrative purposes, data were fitted with an exponential function. Spearman correlation between variables was −0.9 (p = 0.028). (F) Amplitude of phasic response (color) as a function of cumulative O_2_ after each addition, sorted by sensor’s tonic KmO_2_^app^. Amplitudes were calculated as Ch concentration, based on sensors’ response to 5 μM Ch. (G) Maximal response to O_2_ (from each biosensor calibration) as a function of KmO_2_^app^ (n=14). The negative correlation between the variables was significant (r_spearman_ = −0.86, p<0.0001). Data were fitted with an exponential decay curve (fitted decay constant *k* = 0.4 μM^-1^ (CI = 0.031-0.78 μM^-1^, rmse = 0.68 μM). (H) Amplitudes of phasic ChOx responses divided into low and high-KmO_2_^app^ groups (n=7 in each group) following consecutive step additions of O_2_. Control phasic O_2_ values obtained using the same algorithm are also plotted for the same groups (n=6 for high-KmO_2_^app^ and n=4 for high-KmO_2_^app^). Low-KmO_2_^app^ transients were significantly higher than those from the high-KmO_2_^app^ group (p<0.0001) and the amplitudes from both ChOx groups were significantly higher than any O_2_ group (p<0.005). Oxygen transients from low-KmO_2_^app^ vs. high-KmO_2_^app^ groups did not significantly differ (p=0.998). Group comparisons done by two-way ANOVA for unbalanced data followed by Tukey-Kramer post-hoc tests. (I) T50 decay time of transients from low- and high-KmO_2_^app^ groups as a function of cumulative O_2_. The decay of peaks in response to 10 μM cumulative [O_2_] was significantly longer than any other condition (p<0.0005, two-way ANOVA for unbalanced data followed by Tukey-Kramer post-hoc tests). (J) Hill coefficient from the fits of cumulative COA phasic component vs. cumulative O_2_ (as in B) as a function of KmO_2_^app^ (n=14). The two variables were negatively correlated (r_spearman_ = −0.64, p = 0.015), showing an exponential-like relationship (fitted initial amplitude of 1.27 ± 0.75, decay constant of 0.33 ± 0.62 μM^-1^ and offset of 0.97 ± 0.67). *** p<0.001.

Next, we assessed how the coating composition and its physical properties could modulate the sensor O_2_-dependence. First, we calculated the biosensor efficiency (ratio of Choline vs. H_2_O_2_ sensitivities), as a proxy to the enzyme loading in the coating and plotted it against tonic KmO_2_^app^ (Figure 7E). The result revealed a decreasing trend (Figure 7E), suggesting that biosensors with a high enzyme loading have low sensitivity to tonic O_2_ changes (low-KmO_2_^app^). Strikingly, the biosensors with the lowest tonic KmO_2_^app^ exhibited the highest phasic peaks (Figure 7F), with maximal phasic responses to O_2_ decreasing exponentially as a function of KmO_2_^app^ (Figure 7G).

In order to further detail on the effect of O_2_ baseline on non-stationary COA, we split the sensor calibrations into two groups according to their tonic O_2_-dependence. As already suggested in Figure 7D, we confirmed that the larger phasic responses in the low-KmO_2_^app^ vs. high-KmO_2_^app^ groups could not be attributed to differences in the underlying O_2_ profiles. Oxygen transients after each addition were negligible and did not significantly differ across KmO_2_^app^ groups (p=0.998, Figure 7H). Noteworthy, in the low-KmO_2_^app^ group, the highest COA peak was achieved in response to the second O_2_ addition (10 μM of cumulative [O_2_]) rather than to the first in 6 out of 7 calibrations (marginally significant difference between 5 and 10 μM O_2_ responses, p=0.076, paired *t*-test). Furthermore, the same COA peaks had the longest decay across the two KmO_2_^app^ groups (p<0.0005, Figure 7I). These non-monotonic profiles with respect to [O_2_] reflected the deviation of the cumulative phasic COA vs. O_2_ curves from a Michaelis-Menten kinetics. Accordingly, the Hill coefficients extracted from calibration fittings (e. g. Figure 7B) were above 2 for sensors with low KmO_2_^app^ and significantly decreased towards 1 as KmO_2_^app^ increased (p<0.05). These observations suggest a cooperative mechanism that enhances phasic responses as the O_2_ baseline increases, in biosensors with low tonic O_2_ dependence.

In summary, we show that, under a Ch background, ChOx biosensors respond to O_2_ with a transient increase in enzyme activity before reaching a steady-state. Importantly, phasic and tonic components were apparently mutually exclusive and their relative magnitude was sensitive to the properties of the enzyme coating, namely enzyme loading. Sensors with high enzyme loading have a low tonic O_2_-dependence and show large phasic responses that seem to be potentiated by the O_2_ baseline.

A possible explanation for the sensors’ phasic and tonic components and their anti-correlation is the local consumption of Ch within the coating milieu, boosted by phasic O_2_. In that case, a large Ch depletion, in sensors with high enzyme loading, would blunt the tonic ChOx response to O_2_ while sparing the initial non-steady-state peak in enzyme activity.

### Modeling *in vitro* biosensor responses reveals the mechanisms underlying tonic and phasic oxygen-dependence

In order to provide a theoretical ground for our *in vitro* observations, we have simulated the behavior of biosensors in calibration conditions. We numerically solved a system of partial differential equations describing the diffusion of Ch and O_2_ in the coating and their interaction with the enzyme, leading to H_2_O_2_ generation.

To mimic our experimental calibrations, we simulated sensor responses to 5 μM step increases in O_2_ (starting from zero) under a constant level of 5 μM Ch in the bulk solution. Remarkably, in line with our experimental findings, the model predicted phasic and tonic components of sensor response to O_2_ whose magnitude depended on O_2_ baseline (Figure 8A). Sensors with high enzyme loading showed higher phasic peaks and tonic responses that saturate at lower O_2_ baselines than sensors with a low enzyme amount (Figure 8A). To get a more resolved characterization of biosensor’s O_2_ dependence, we next generated response curves at 1 μM O_2_ steps, until saturation of enzymatic H_2_O_2_ generation was nearly reached (see Methods). By simulating a range of coating thicknesses and enzyme concentrations, we found that both parameters decreased KmO_2_^app^ of tonic responses (Figure 8B) and increased the magnitude of phasic peaks (Figure 8C). Interestingly, our model predicts that particular combinations of coating thickness and enzyme concentration can be used to optimize sensor sensitivity to Ch (at saturating O_2_ levels) (Figure S6A). Yet, such a strategy is not expected to concomitantly reduce phasic and tonic O_2_ dependence, which seem to be mutually exclusive. In agreement with the experimental data, our model anticipated that sensors with the lowest KmO_2_^app^ exhibit the highest transient responses to O_2_ (Figure 8B-D). Across the simulated coatings, the highest phasic peaks occurred when sensor cumulative tonic responses were close to maximal and progressively decreased in sensor tonic saturation level as a function of tonic KmO_2_^app^ (Figure 8D). This observation is in agreement with our experimental observation of highest phasic peaks at a 10 μM O_2_ baseline (Figure 7H-J).

**Figure 8.**
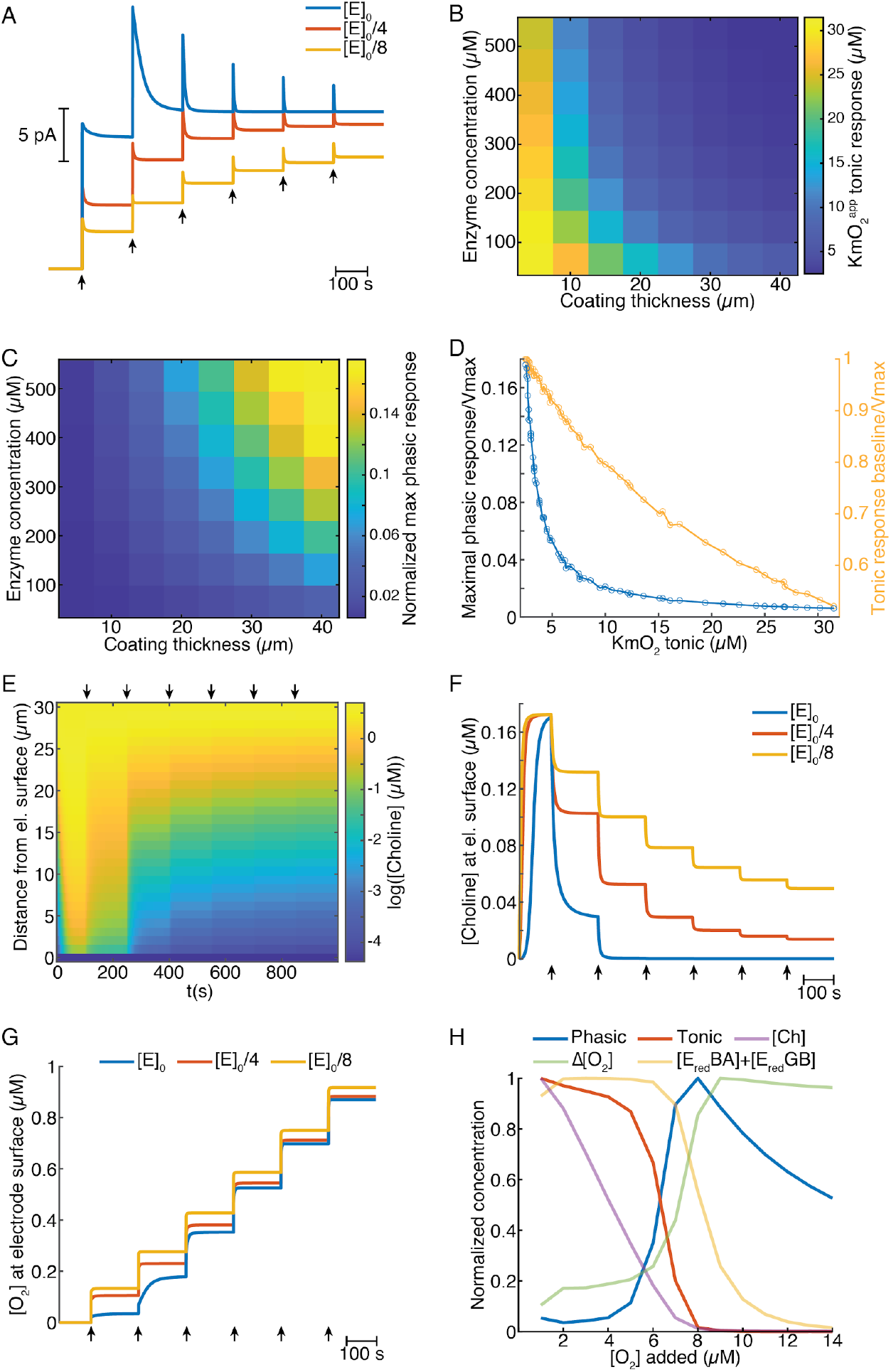
Mathematical model explains ChOx-based biosensor COA responses to oxygen. (A) Simulated calibrations of biosensors with different enzyme concentrations in the coating (coating thickness of 30 μm). Choline in bulk solution was kept constant at 5 μM and O_2_ was incremented in 5 μM steps from zero to 30 μM at times indicated by arrows. (B) Apparent KmO_2_ of tonic sensor responses as a function of coating thickness and enzyme concentration in the coating. (C) Normalized maximal phasic responses of sensors with different coating thicknesses and enzyme concentrations. Phasic component magnitudes for each sensor refer to the highest phasic response divided by the maximal cumulative tonic response (Imax) from the respective simulated sensor calibration. (D) Blue trace represents the normalized maximal phasic response vs. tonic KmO_2_^app^ and the orange trace refers to the level of saturation of the sensor’s tonic response at which the maximal phasic peak occurs. Data were obtained from all combinations of coating thickness and enzyme concentration in B and C. (E) Concentration profile of Ch in the sensor coating as a function of distance from the electrode surface during a simulated calibration of a high enzyme-loaded sensor (coating thickness of 30 μm and enzyme concentration of 560 μM, same as the blue trace in A). Arrows indicate 5 μM O_2_ step increments in solution. (F) Time-course of Ch concentration at the electrode surface of sensors with different enzyme concentrations during simulated calibrations. Arrows indicate 5 μM O_2_ step increments in solution. (G) Timecourse of O_2_ concentration at the electrode surface for sensors with different enzyme concentrations during simulated calibrations. Arrows indicate 5 μM O_2_ step increments in solution. (H) Normalized profiles of phasic and tonic sensor responses as well as of concentrations of enzyme substrates and total reduced enzyme-bound complexes at the electrode surface as a function of O_2_ in solution. Data are from a simulated calibration of a sensor with a 30 μm coating and an enzyme concentration of 560 μM, upon 1 μM O_2_ step increases in solution. The ΔO_2_ is the initial rise in O_2_ following each O_2_ increment in solution (at a lag of 0.3 s).

To get further clues into the factors shaping sensors’ O_2_-evoked responses, we analyzed the concentration dynamics of Ch and O_2_ in the coatings during simulated calibrations. We observed that, under high enzyme loading, Ch is rapidly depleted in the coating as O_2_ levels in solution increase (Figure 8E). This effect is less pronounced in sensors with low enzyme loading (Figure 8F). Interestingly, significant O_2_ consumption, observed mainly in coatings highly loaded with enzyme, was stronger for low O_2_ levels before reaching saturation of the sensor tonic response (Figure 8G). These observations suggest that depletion of Ch in the enzyme coating is the limiting factor that shapes sensors’ tonic responses to O_2_.

To further investigate phasic responses under high enzyme loading, in addition to substrate profiles, we assessed the levels enzyme-bound reduced intermediate species vs. buffer O_2_ concentration, as those are direct precursors of H_2_O_2_. We observed that, under non-saturating O_2_ levels, the concentrations of EredBA and EredGB change oppositely as O_2_ is increased, which maintains the sum of both intermediates relatively stable (Figure S6B). At 1 μM buffer O_2_ steps, the profiles of reduced intermediates, Ch and O_2_ at the electrode surface, suggest that phasic biosensor peaks result from a combination of multiple factors (Figure 8H). As expected, the sensor’s tonic response steeply decays upon Ch depletion in the coating. This decrease is accompanied by a sharp rise in the instantaneous ΔO_2_ at the electrode surface evoked by O_2_ increments in the bulk solution. As the rate of O_2_ consumption depends on the concentration of reduced enzyme-bound complexes, the increasing profile of ΔO_2_ results, in a first stage, from the depletion of the EredBA complex, followed by the decrease in E_red_GB (Figure S6B). In turn, the amount of E_red_BA and E_red_GB depends on both Ch and O_2_, which leads to a summed profile that is shifted to the right relative to the sensor’s tonic response. Thus, phasic sensor peaks reach maximal levels at the offset of tonic responses, when there is a combination of relatively large concentrations of the direct H_2_O_2_ precursors, namely enzyme-bound reduced complexes and O_2_, and very low levels of Ch in the coating (Figure 8H).

Overall, our biosensor simulations corroborate the *in vitro* results, firstly converging towards the notion that Ch consumption in the coating is a major factor governing sensor tonic O_2_ dependence. Secondly, the results suggest that non-steady state phasic responses depend on the relative concentrations of Ch, reduced intermediate enzyme complexes and O_2_ in the coating.

## Discussion

We have developed a novel multi-site tetrode-based amperometric ChOx (TACO) sensor optimized for the highly sensitive and unbiased simultaneous measurement of putative cholinergic activity and O_2_ dynamics in the brain. Our approach, based on the differential plating of recording sites to create *pseudo-sentinel* channels outperforms previous common-mode rejection strategies, which were limited by diffusional cross-talk (Burmeister et al., 2003; Parikh et al., 2007; Santos et al., 2015). This strategy allowed us to substantially reduce the size and increase the spatial confinement of recording sites by using a 17 μm wire tetrode as the biosensor electrode support. Our recordings in freely-moving and head-fixed rodents reveal the usefulness of this compact multi-site design to clean artifacts from sensor signals and assess the correlation between the activity of the immobilized enzyme and brain extracellular O_2_ on a fast time-scale. Importantly, this method can be generalized to improve the selectivity and address the *in vivo* O_2_-dependence of any oxidase-based biosensor.

By simultaneously measuring the activity of immobilized ChOx and extracellular O_2_ in the hippocampus of behaving mice, we found that fast biosensor signals correlate in amplitude and time with O_2_ transients evoked by behavioral and network dynamics events exemplified by locomotion bouts and SWRs. Notably, the relationship between COA and O_2_ profiles was apparently not sensitive to the underlying neurophysiological or behavioral context and was preserved during periods without appreciable SWR incidence or locomotion. By using two different methods to manipulate extracellular O_2_, we show that O_2_ fluctuations in the physiological range can drive phasic COA. Remarkably, the time-scale dependence of biosensor response amplitude and lag relative to exogenous O_2_ qualitatively matched that of spontaneous profiles, suggesting that the same directionality happens in spontaneous conditions.

Locomotion-related O_2_ elevations in head-fixed mice have been recently shown to be modulated mainly by respiration rate (Zhang et al., 2019), whereas SWR-evoked O_2_ peaks have been indirectly inferred by fMRI and likely result from neurovascular coupling (Ramirez-Villegas et al., 2015). Thus our study provides a link between the neurophysiological or systemic mechanisms that modulate brain O_2_ levels and the response of ChOx-based biosensors *in vivo*. As it is an intrinsic component of any behavioral task, our results highlight the importance of controlling for O_2_-evoked biosensor signals related to locomotion or movement. It is thus likely that, in reward-related tasks, locomotion related to reward retrieval elicits few seconds delayed phasic changes in O_2_ that drive transient increases in COA. Likewise, a high incidence of SWRs in reward locations, reflecting a consummately state (Buzsáki, 2015), might trigger O_2_ transients and, in turn, phasic ChOx biosensor responses. These two examples provide alternative explanations for previously reported cholinergic transients inferred from COA signals in the prefrontal cortex and hippocampus of rodents engaged in cognitive tasks (Howe et al., 2017; Parikh et al., 2007; Teles-Grilo Ruivo et al., 2017). Importantly, since the rate of the enzymatic reaction depends on both substrates, biosensor responses caused by O_2_ transients are expected to decrease following experimental controls that have been aimed at inhibiting or removing cholinergic inputs (Parikh et al., 2007).

Our *in vitro* characterization of the biosensor O_2_ dependence provided critical insights to interpret the *in vivo* relationship between COA and O_2_. We found robust O_2_-evoked phasic responses whose amplitude was anti-correlated with sensors’ tonic O_2_ dependence. Interestingly the phasic peaks decreased with O_2_ baseline but were detected even under relatively high O_2_ levels, suggesting a high likelihood of such responses to occur *in vivo.* Thus, our data emphasize the impact of O_2_-evoked nonsteady-state biosensor dynamics on fast time-scale *in vivo* measurements. This has not been described in previous studies partly because the experimental procedures used to study O_2_-dependence were unable to unmix tonic and phasic components of sensor response (Baker et al., 2017; Burmeister et al., 2003; Dixon et al., 2002). Instead of generating a continuous O_2_ increase, we induced step increments in O_2_, allowing temporal deconvolution of sensor response components and unbiased estimation of the corresponding K_0.5_O_2_^app^ values.

Remarkably, most of our observations related to sensor O_2_-dependence were predicted by mathematical simulation of biosensor responses *in vitro*. The model incorporated the kinetics of the enzyme-catalyzed reaction, including enzyme-bound intermediate species, and simulated substrates’ diffusion into the coating and interaction with the enzyme leading to H_2_O_2_ formation. In agreement with the experimental data, the model predicted that tonic and phasic components of O_2_ dependence are mutually exclusive. Furthermore, our simulations suggest that increasing the enzyme loading, either by changing enzyme concentration or the coating thickness, amplifies sensor phasic responses and reduces tonic KmO_2_^app^. It is therefore apparent that there is no perfect combination of these two parameters that can concomitantly minimize the two components of biosensor O_2_ dependence. In addition to reinforcing our experimental conclusions, the biosensor model provided important insights into the factors that determine phasic and tonic components of O_2_ dependence. Simulated profiles showed that tonic responses to O_2_ are largely shaped by Ch depletion in the coating, which limits the linearity of the sensor response. On the other hand, phasic responses depended on the instantaneous balance between the concentration of substrates and reduced enzyme-bound intermediates, which lead to the formation of H_2_O_2_. Simulated transients were highest when the concentration of reduced intermediates and O_2_ were in relative excess in comparison to Ch. Since variations in enzyme concentrations, coating thickness and diffusional components have mainly a quantitative effect in our estimates, these conclusions can be extrapolated to sensor geometries and enzyme coating compositions that we have not covered. Furthermore, mathematical model of the biosensor described here provides a rigorous approach for exploration and optimization of the design of any future enzymatic biosensors and investigation of their behavior under non-stationary in vivo conditions.

The phasic O_2_-evoked COA signals described *in vitro* and predicted by mathematical modeling provide crucial information to interpret the time-scale dependence of *in vivo* sensor dynamics triggered either to spontaneous or exogenously evoked O_2_ peaks. In light of those results, the temporal advancement and amplitude drop of biosensor transients relative to O_2_, as O_2_ rises for longer periods, is compatible with a major contribution of the phasic component of biosensor’s O_2_-dependence. This observation suggests that, *in vivo,* our biosensors operated in a regime close to saturation of the tonic response and highlight the effect of non-steady-state ChOx biosensor responses to O_2_, which can be erroneously attributed to Ch.

Our observations disfavor the quantitative optimization of coating properties as a strategy to reduce sensors’ O_2_ dependence. Instead, we anticipate that strategies that increase O_2_ accumulation in the enzyme coating (Njagi et al., 2008) might have relative success, although the high O_2_ levels required to cancel phasic responses are hard to reach passively. Alternatively, some oxidase-based biosensors for *in vitro* applications have incorporated an electrochemical actuator that enables local manipulation of O_2_ concentration based on water electrolysis (Park et al., 2006). However, applying such design *in vivo* would require miniaturization and separation between the O_2_ generation compartment and the brain to avoid electrolytic tissue damage. Furthermore, in addition to the main O_2_ confound, one cannot completely rule out a possible modulation of COA *in vivo* by factors that affect enzyme conformation, including temperature and pH (Hekmat et al., 2008). Although the enzyme is poorly sensitive to the modest variations of these factors *in vivo* (Csernai et al., 2019; Venton et al., 2003), it would be relevant to characterize their potential contribution to COA dynamics in future studies. A further validation of ChOx-based measurements could be achieved by confronting the dynamics of COA with that of cholinergic signals measured with other sensing approaches, under the same experimental conditions. The latter technics include optogenetically-taged single unit recordings or fluorescence reporters, which have previously revealed fast cholinergic dynamics related to arousal, sensory sampling, negative reinforcements and unexpected events (Eggermann et al., 2014; Hangya et al., 2015; Lovett-Barron et al., 2014; Reimer et al., 2016).

In summary, our results suggest that ChOx biosensor signals *in vivo* are composed of a mixture of O_2_-related artifacts and true cholinergic dynamics. The weight of each factor depends on the time-scale, with slow state-related changes reflecting cholinergic dynamics with low O_2_-related contamination and fast transients, except for a minor fraction (less than ca. 5% in our data), being caused by phasic O_2_ fluctuations. We show that O_2_ transients can be triggered by cognitively-relevant events, such as locomotion and periods of high SWR incidence, and confound the ChOx-based measurement of cholinergic activity. Thus, our study reveals a previously ignored phasic O_2_-dependence of ChOx-based biosensors which is critical to control for, particularly in the case of fine time-scale measurements of ACh.

Importantly, our conclusions can probably be generalized to other oxidase-based biosensors that have been used to measure neurotransmitters or metabolically-relevant molecules in the brain (Chatard et al., 2018; Dixon et al., 2002; Hascup et al., 2013; McMahon et al., 2007). The exact extent of phasic and tonic O_2_ dependence would depend on the particular enzyme kinetics and on the basal extracellular concentrations of analyte relative to the magnitude of changes in the brain.

## Materials and Methods

### Chemicals and solutions

All chemicals were of analytical grade, purchased from Sigma-Aldrich and used as received. Solutions were prepared in ultra-pure deionized water (≥18MΩ.cm) from a Milli-Q water purification system.

### Tetrode fabrication and platings

The microelectrode support material was a 17 μm diameter Platinum/Iridium (90/10) wire insulated by a polyimide coating (California Fine Wire Company). Tetrodes were fabricated using standard methods (Gray et al., 1995). Briefly, four wires were twisted together and heated to melt the insulation, creating a stiff bundle of twisted wires. The wires’ insulation at the untwisted ending of the tetrode was then removed and the tetrode was inserted in a silica tube (150 μm inner diameter), which was glued to a holder that allowed easy manipulation of the tetrode. Next, the untwisted endings of the tetrode wires were soldered to the pins of an adapter fixed to the tetrode holder, allowing connection to the potentiostat’s head-stage. Finally, the twisted ending of the tetrode was cut using micro-serrated stainless-steel scissors.

Tetrode surface treatments and platings were performed with a portable potentiostat (EmStat 3, PalmSens BV), using a freshly-prepared Ag/AgCl wire (125 μm diameter, WPI inc.) as *pseudo*reference electrode. Prior to platings, electrode surfaces were cleaned by swirling the tetrode tip in isopropanol followed by an electrochemical treatment in PBS. For that purpose, we applied 70 cycles of a square wave with a first step at +1.2 V for 20 seconds followed by a 4 seconds step at −0.7 V. All tetrode sites were then gold-plated in a 3.76 μM aqueous solution of tetrachloroauric acid by applying 20 cycles of a square wave that alternated between +0.6 V for 10 s and −1.0 V for 10s. Sites’ impedances were checked after gold-plating using a nanoZ impedance tester (Multichannel Systems, GmbH). The pair of sites with the highest impedance was then platinized in a 10 mM chloroplatinic acid solution in 0.1 M sulfuric acid by DC amperometry at −0.1 V, until a current of −30 nA was reached.

### TACO sensor coatings

Choline oxidase immobilization was performed as previously described (Santos et al., 2015). Briefly, a 0.5% (w/v) chitosan stock solution was solubilized in saline (0.9% NaCl) under stirring at pH 4-5, adjusted by addition of HCl. After solubilisation, the pH was set to 5-5.6 by stepwise addition of NaOH.

To form a chitosan/ChOx cross-linked matrix, 1.5 mg of p-benzoquinone was added to 100 μL of 0.2% chitosan. Four μL of this solution were then mixed with a 4 μL aliquot of ChOx at 50 mg/mL in saline. The tetrode tip was coated by multiple dips (10-15) in a small drop of ChOx immobilization mixture, created using a microliter syringe (Hamilton Co.). The microelectrode and syringe were micromanipulated under a stereomicroscope. The coating procedure was stopped when the chitosan/protein matrix was clearly visible under the microscope.

Following enzyme immobilization, tetrode site’s response to Ch was tested and *meta*phenylenediamine (*m*-PD) was electropolymerized on the pair of sites with the highest sensitivity (please see *Biosensor calibrations* sub-section). Electropolymerization was performed in a nitrogen bubbled oxygen-free PBS solution of 5 mM *m*-PD by DC amperometry at +0.6 V during 1500 s. The biosensors were stored in air and calibrated on the day after *m*-PD electropolymerization.

### Biosensor calibrations

All *in vitro* tests were done in a stirred calibration buffer kept at 37 °C using a circulating water pump (Gaymar heating/cooling pump, Braintree Scientific, Inc., USA) connected to the calibration beaker. Routine calibrations after enzyme immobilization and *m*-PD electropolymerization steps were performed by amperometry at a DC potential of +0.6 V vs. Ag/AgCl *pseudo-reference* electrode. After stabilization of background current in PBS, sensors were calibrated by three consecutive additions of 10 μM Ch followed by 4.9 μM H_2_O_2_. In the case of complete (*m*-PD polymerized) biosensors, the response to 1 μM DA and 100 μM AA was also tested. Voltammograms of H_2_O_2_ were done by consecutive additions of 4.9 μM H_2_O_2_ at different applied DC voltages.

*In vitro* O_2_ tests were carried out in a sealed beaker. After addition of 5 μM Ch, the calibration buffer was bubbled with nitrogen during approximately 30 min. Then, known O_2_ concentrations (5 or 10 μM) were added to the medium from an O_2_-saturated PBS solution previously bubbled with pure O_2_ during 20 min. Biosensor response to O_2_ additions was measured at +0.6 V *vs.* Ag/AgCl. In a set of calibrations, O_2_ was concurrently measured by polarizing a gold-plated channel at −0.2 V, while keeping the remaining at +0.6 V. To obtain O_2_ voltammograms, calibrations consisting in 3 additions of 5 μM O_2_ were performed at different applied DC potentials. In these experiments, O_2_ was purged from the solution after each voltage step.

### Experimental model and subject details

Freely-moving recordings were performed on a 6 months old Long-Evans rat and head-fixed recordings were done in a total of six 3-7 month old C57BL/6J mice. All experimental procedures were established, and have been approved in accordance with the stipulations of the German animal welfare law (Tierschutzgesetz)(ROB-55.2-2532.Vet_02-16-170).

### Surgeries

For freely-moving recordings, we chronically implanted a tetrode biosensor and a 32-channel linear silicon probe array (A1×32-7mm-100-1250-H32, NeuroNexus Technologies, Inc) in the rat brain. The general procedures for chronic implantations of electrode arrays have been described in detail (Vandecasteele et al., 2012). Prior to surgery, the tetrode biosensor and the silicon probe array were attached to home-made microdrives. Silicon probe’s sites were then gold-plated until impedances at 1 kHz decreased below 200 kΩ (Ferguson et al., 2009). Anesthesia was induced with a mixture of Fentanyl 0.005 mg/kg, Midazolam 2 mg/kg and Medetomidine 0.15 mg/kg (MMF), administered intramuscularly. The rat was continuously monitored for the depth of anesthesia (MouseStat, Kent Scientific Corporation, Inc.). After the MMF effect washed out, anesthesia was maintained with 0.5-2% isoflurane via a mask, and metamizol was then subcutaneously administered (110 mg/kg) for analgesia. The tetrode biosensor was implanted in the cortex above the right dorsal hippocampus (AP −3.7 mm, ML −2.5 mm, DV −1.2 mm, relative to bregma) and the silicon probe array was implanted at 0.8 mm posterior from it, spanning most cortical and hippocampal layers (AP −4.5, ML −2.4, DV – 3.4). The microdrives were secured to the skull with a prosthetic resin (Paladur, Kulzer GmbH). An Ag/AgCl (125 μm thick) silver wire coated with Nafion (Hashemi et al., 2011) was inserted in the cerebellum and served as the *pseudo*-reference electrode for electrochemical recordings. The ground for electrophysiology was a stainless-steel screw implanted at the surface of the cerebellum. To reduce line noise, this Ag/AgCl wire was shorted with the electrophysiology ground at the input of the electrochemical head-stage.

Mice used in head-fixed recordings were implanted with a head-post. Anesthesia followed the same procedures as in rats. A mixture of 0.05 mg/kg Fentanyl, 5 mg/kg Midazolam and 0.5 mg/kg Medetomidine was administered intraperitoneally to induce anesthesia, which was later maintained with isoflurane and Metamizol (200 mg/kg). A craniotomy was made above the dorsal hippocampus and a Nafion-coated Ag/AgCl wire was implanted in the cerebellum. Depending on the head-post configuration, it was cemented either to the back of the skull above the cerebellum or above the hemisphere contralateral to the craniotomy, using UV-curing dental cement (Tetric EvoFlow, Ivoclar Vivadent AG). Finally, the craniotomy and surrounding skull were covered with a silicone elastomer (KWIK-CAST, World Precision Instruments Inc.).

### Electrochemical and electrophysiological equipment and recordings

Amperometric measurements were performed using either a 4-channel (MHS-BR4-VA) or a 8-channel (MBR08-VA) potentiostat connected to 4- or 8-channel miniature head-stages, respectively (npi electronic GmbH, Germany). In addition to providing a higher channel count, the MBR08-VA allowed independent control of the potential applied to each channel. This feature enabled simultaneous measurement of the biosensor signal, arising from ChOx, and O_2_. The DC analog signal from the head-stage was amplified and digitized at 30 kHz and stored for offline processing using the Open Ephys acquisition board and GUI (Siegle et al., 2017).

Freely-moving electrochemical recordings were done using the MHS-BR4-VA potentiostat and the corresponding 4-channel miniature head-stage. Electrophysiological signals were pre-amplified using a 32-channel head-stage with 20x gain (HST/32V-G20, Plexon Inc.) which was connected to a multichannel acquisition system (Neuralynx Inc). Data was acquired at 32 kHz and stored for offline processing. Both head-stages were connected to the respective recording systems via light and flexible cables suspended on a pulley so as not to add weight to the animal’s head. The tetrode biosensor was gradually lowered through the cortex until it reached the hippocampal CA1 pyramidal layer. Correct targeting was assessed based on brain atlas coordinates and by the identification of hippocampal ripples. Recordings were performed in a square open-field arena (1.5 m x 1.5 m), where the animal could sleep or explore the environment at will. Chocolate sprinkles were occasionally spread on the maze to enforce exploratory behavior. The position of the rat head was derived from small reflective markers attached to the chronic implant. A motion capture system consisting of multiple infrared cameras (Optitrack, NaturalPoint Inc.) was used to 3D-track the markers with high spatio-temporal resolution (data acquired at 120 Hz).

Head-fixed recordings in mice were performed using the MBR08-VA potentiostat and respective head-stage. After fixing the mouse, the layer of silicone elastomer protecting the craniotomy was removed. The dura matter above the target brain region was removed and the tetrode biosensor was slowly inserted through the cortex until the hippocampal CA1 pyramidal layer was reached. Accurate targeting was assessed according to brain atlas coordinates and/or by the online identification of hippocampal ripples in the recording. In 5 out of 10 recording sessions mice were head-fixed on a cylindrical treadmill. Movement was quantified based on the video optical flow arising from treadmill rotation using Bonsai (Lopes et al., 2015). In the remaining sessions, mice were head-fixed on a rotating disc which encoded its turns. The analog signal from the disc encoder was fed into the Open Ephys acquisition board and used to quantify mice locomotion.

### Data analysis

Raw recordings were preprocessed by low-pass filtering and resampling at 1 kHz. All data analysis was done in Matlab using custom-made functions (MathWorks).

#### *In vitro* sensor responses

*In vitro* analysis of biosensor responses was performed on 10 Hz downsampled data, low-pass filtered at 1 Hz. Sensitivities to Ch and O_2_ were determined by linear-regression of the responses to the first 3 analyte additions, whereas the sensitivities to H_2_O_2_ and interferents were estimated from a single addition. Minimal selectivity ratios were estimated by dividing the lower limit of the 95% confidence interval (CI) of Ch sensitivity by the upper limit of the CI of interferent responses. Following *pseudo*sentinel subtraction, the biosensors’ limit of detection (LOD) for Ch was calculated as the Ch concentration corresponding to 3 times the baseline standard deviation (SD). The T50 and T90 response times were defined as the time between the onset of current increase in response analyte and 50% or 90% of the maximum current, respectively.

#### Artifact cancellation by common-mode rejection

*In vivo* electrochemical signals from Ch-or O_2_-sensitive sites were cleaned by subtraction of the respective 1 kHz data by the *pseudo*-sentinel (Au-Pt/*m*-PD) channel upon a frequency-domain correction. The latter procedure has been described in detail and optimizes common-mode rejection by correcting phase and amplitude mismatches between channels arising from slight variations in impedance (Santos et al., 2015). This correction was based on the estimation of a transfer coefficient *(T)* describing the transfer of LFP-currents from Sentinel to the Ch-sensitive (ChSens) channel in the complex domain, according to the following equation:

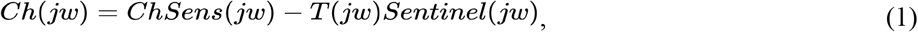

where *jw* is a complex value at frequency *w* and *Ch* is the clean putative Ch signal. Upon applying a fast Fourier transformation (FFT) on both signals, the amplitude of *T* for each FFT frequency bin was estimated from the square root of ratio between the power of ChSens and sentinel channels. The phase of T was estimated from the phase of the ChSens/sentinel cross-spectrum (phase-shift) during timewindows with high phase-locking (>0.98). These estimations were computed for each biosensor used *in vivo* from average spectra obtained from multiple slow-wave periods. In the low frequency range (<0.3 Hz), due Ch contribution to power in the ChSens channel, the estimation of the amplitude of *T* was done by linear extrapolation considering the trend at contiguous higher frequencies. The cleaned signal of putative *Ch* was obtained by inverse FFT of *Ch*(*jw*). Cleaned O_2_ signals were obtained following the same logic.

Clean Ch and O_2_ signals were then low-pass filtered at 1 Hz and downsampled to 10 Hz for most of the analysis excluding time lags, which were computed on 100 Hz downsampled data.

#### Brain state separation

Local-field potential-related power spectrograms were computed using custom-made Matlab functions based on multi-taper analysis methods (Mitra and Pesaran, 1999). Separation of brain states in freely-moving recordings was based on LFP spectral features and behavior. Active wake states were defined as periods when the animal moved vigorously and continuously (>30 s) and showed a prominent LFP spectral peak in theta range (6-10 Hz). Quiet wakefulness or immobility was defined as a period without prominent theta and with occasional movements (<30 s between movement bouts). Long periods (>1 min) without movement and without prominent theta, rather showing high delta power (1-4 Hz) were ascribed to NREM sleep. Rapid eye movement sleep was detected as periods showing a sustained theta band (>30 s) and negligible movement.

#### Sharp-wave/ripples and related biosensor signals

To detect SWRs, the wide-band electrochemical or electrophysiological signal was band-pass filtered (120-200 Hz), squared and smoothed with a 4.2 ms standard deviation and 42 ms wide Gaussian kernel. The square root of this trace was then used as the power envelope to detect oscillatory bursts. The events exceeding the 98^th^ percentile of the power envelope distribution, having at least 5 cycles and lasting less than 200 ms were detected as ripples. For the analysis of correlations between integrated ripple power and SWR-triggered sensor signals, a ripple power envelope was obtained upon Hilbert-transforming the band-pass filtered electrochemical signal (ripple band, 120-200 Hz). Different ripple integration times were obtained by smoothing the power envelope with moving average windows of different lengths. Correlations were then computed between smoothed ripple power envelopes at SWR times and the corresponding changes in COA or O_2_ (relative to their baseline value 1s prior to SWRs) at different lags from SWRs.

The amplitudes of SWR-related COA and O_2_ were obtained from the difference between the values at SWR lags corresponding to peaks and onsets, extracted from average SWR-triggered traces.

#### Locomotion bouts

To detect locomotion bouts in freely-moving recordings, speed was computed from the derivative of low-pass filtered position (0.5 Hz). Speed was then band-pass filtered (0.02-0.2 Hz) and locomotion bouts were detected as peaks in speed that exceeded a manually-set threshold. In head-fixed recordings on the disc, mouse locomotion was derived from its rotation in 1 s bins. When head-fixed on the treadmill, mouse locomotion was quantified based on the optic flow from a recorded video, choosing a region of interest that covered only a moving part of the treadmill. The signal was resampled to 1 Hz in order to match the sampling rate of locomotion on the rotating disc. Locomotion bouts were detected as peaks on the band-pass filtered speed (0.02-0.2 Hz) that exceeded a manually-defined threshold.

The amplitude of locomotion-bout-associated COA and O_2_ signal transients was calculated based on the difference between the values at manually-determined times of transient onsets and peaks, associated with each event. Likewise, the times of locomotion bout onsets (used to calculate speed change) and the associated peaks in theta power were manually defined based on visual inspection of speed time courses and LFP spectrograms, respectively.

#### Oxygen-related ChOx transient signals analysis

The amplitude of broad-band spontaneous and exogenously-induced O_2_ transients and COA transients associated with them was calculated, for each event, as the difference between the peak value and that at semi-automatically defined time of the transient’s onset.

To capture and separately analyze spontaneous COA and O_2_ transients of highly variable timescale (1-14 seconds) and non-harmonic shape we performed peak detection on the two signals band-pass filtered in a frequency range [*f/2 f* with corner frequency *f* defining each filter, varying from 0.05 to 0.5 Hz. Peaks of the transients were then detected in both band-passed COA and O_2_ signals, selecting the events that exceeded half of the maximal peak amplitude. In each frequency band, COA transients used for the analysis were restricted to be within *0.75/f* of any O_2_ peak. The amplitudes of O_2_ and COA transients were defined as peak magnitudes derived from the band-pass filtered signals. For each filter band, the linear relationship between COA and O_2_ peak amplitudes was estimated as a slope of the linear model fit to all detected event pairs for this filter. To interpret the nonlinear shape of the O_2_ transients captured by each filter, we extracted rise time (from trough to peak) from the median peak aligned O_2_ transients and used these values in Figures 4E-H.

The transient onsets and peaks of exogenously-induced changes in O_2_ and CAO were manually detected.

#### Modeling biosensor responses *in vitro*

We simulated biosensor responses *in vitro* by numerically solving a system of partial differential equations that describe the diffusion of the substrates Ch and O_2_ in the enzyme coating and their interaction with the enzyme, leading to product formation.

The buffer solution where the biosensor was placed for calibration is a free-flow environment, in which the concentrations of Ch and O_2_ are constant over time. Therefore, considering *R* is the coating thickness, at the boundary between the biosensor coating and the calibration buffer Ch is kept constant at 5 μM during the calibration.

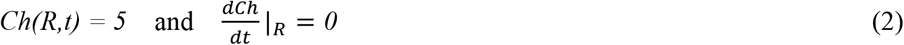

Oxygen is changed in steps (*x*), starting from 0, during the calibration, so between O_2_ step increases we have:

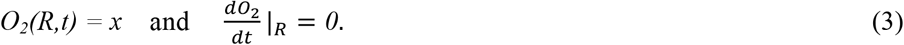

Enzyme substrates (Ch and O_2_) diffuse from the bulk solution into the enzyme layer, eventually reaching the electrode site, whereas H_2_O_2_ is locally generated and diffuses within the coating. As the size of our recording sites is very small, this process is better described by a spherical diffusion equation:

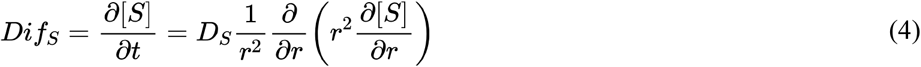

where *S* represents substrates or H_2_O_2_ concentration, *D_S_* is the respective diffusion coefficient and *r* is the distance to the electrode surface.

In order to simulate realistic two-substrate biosensor responses, we modeled the formation of enzyme intermediate complexes resulting from Ch binding to the enzyme and O_2_ oxidation reactions, which have been described in detail (Fan and Gadda, 2005) (Scheme 1). Briefly, enzyme-bound Ch *(ECh),* which is in equilibrium with the free reactant species (*E* and *Ch*), undergoes a chemical step leading to the reduction of the FAD enzyme prosthetic group. In this nearly irreversible step, Ch is converted to betaine aldehyde, which remains mostly enzyme-bound (*E*_red_BA). The first step in which H_2_O_2_ is produced results from the oxidation of FAD_red_ by O_2_ (*E_ox_BA*) followed by a second chemical step in which FAD_ox_ is reduced by betaine aldehyde. The resulting enzyme-bound glycine betaine (*E_red_GB*) is then oxidized by O_2_, producing H_2_O_2_. The reaction cycle is completed with release of glycine betaine bound to FAD-oxidized enzyme (*E_ox_GB*).

The instantaneous change in the concentration of enzyme substrates, free enzyme, enzyme-bound intermediate complexes and reaction products can then be described by the following system of partial differential equations:

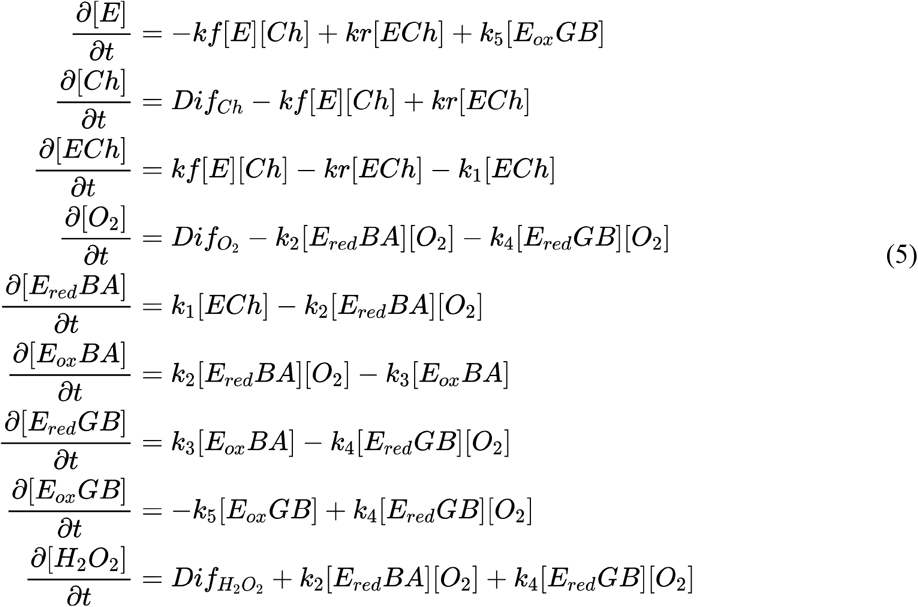

Note that we ignored the kinetics of glycine betaine formation, as it is not relevant regarding the sensor signal transduction and it would not affect the concentrations of any reaction species.

In the beginning of the simulated calibration, there is no Ch in the coating, so all enzyme molecules are free, at a concentration that equals the total enzyme concentration *([E]_0_)*.

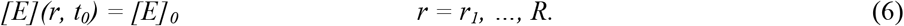

In turn, in the same conditions, there are no enzyme-bound species and no product formed.

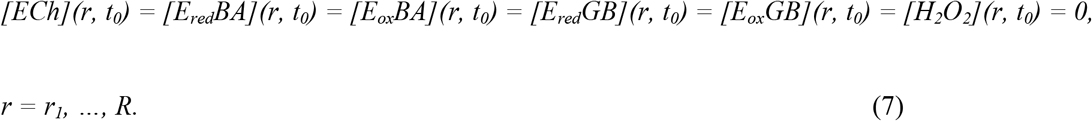

At the electrode surface there is zero flux of substrates and H_2_O_2_ is rapidly oxidized, whereas at the boundary between the coating and bulk solution it is rapidly washed away.

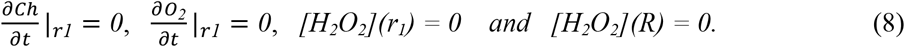

The sensor response current is then given by the charge of two-electrons per each oxidized H_2_O_2_ molecule times the flux of H_2_O_2_ at the electrode surface.

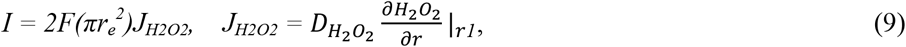

where *F* is the Faraday’s constant and *πr_e_^2^* is the electrode surface area.

Considering the initial conditions described above the system of partial differential equations was numerically solved by discretization in space and time (time and space steps *dt* = 0.1 ms and *dr* = 1 μm, respectively) using the finite difference approximation method (Baronas et al., 2009).

The values ascribed to the variables used in the model are summarized in Table 2. The rate constants corresponding to the reaction mechanism in Scheme 1 were extracted from the corresponding literature (Fan and Gadda, 2005). An enzyme concentration in the coating of 263 uM was estimated from its concentration in the mixture used for coatings, ignoring drying effects upon coating. Variations in enzyme concentration were simulated around that value. We estimated the diffusion coefficients of enzyme substrates and H_2_O_2_ in the coating taking into account the free diffusion coefficients in solution multiplied by a hindrance factor α. The latter was set at 0.8, considering the expected effect of macromolecular crowding on diffusion, for protein concentrations in the range of those used in our simulations (Lamers-Lemmers et al., 2000; Santos et al., 2011)

**Scheme 1.**
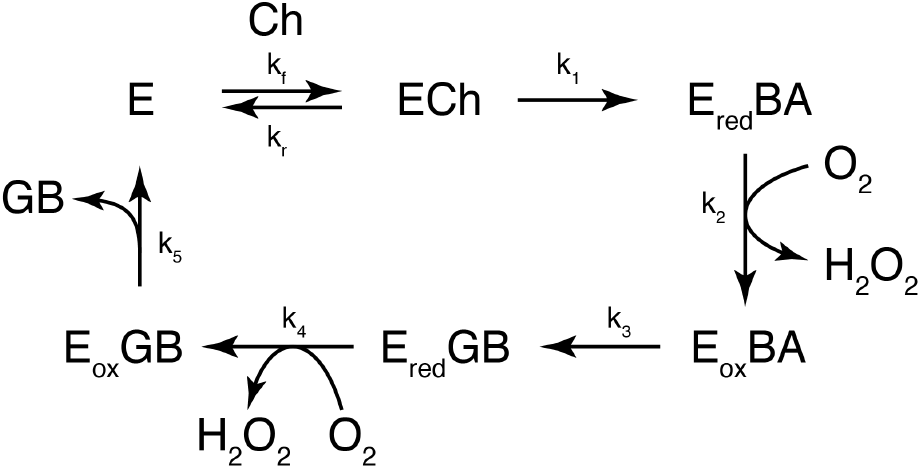
Minimal kinetic mechanism of Choline Oxidase. (E) Free enzyme, (ECh) enzyme-bound Ch, (E_red_) enzyme with reduced FAD co-factor, (E_ox_) enzyme with oxidized FAD co-factor, (BA) betaine aldehyde, (GB) glycine betaine.

**Table 2.**
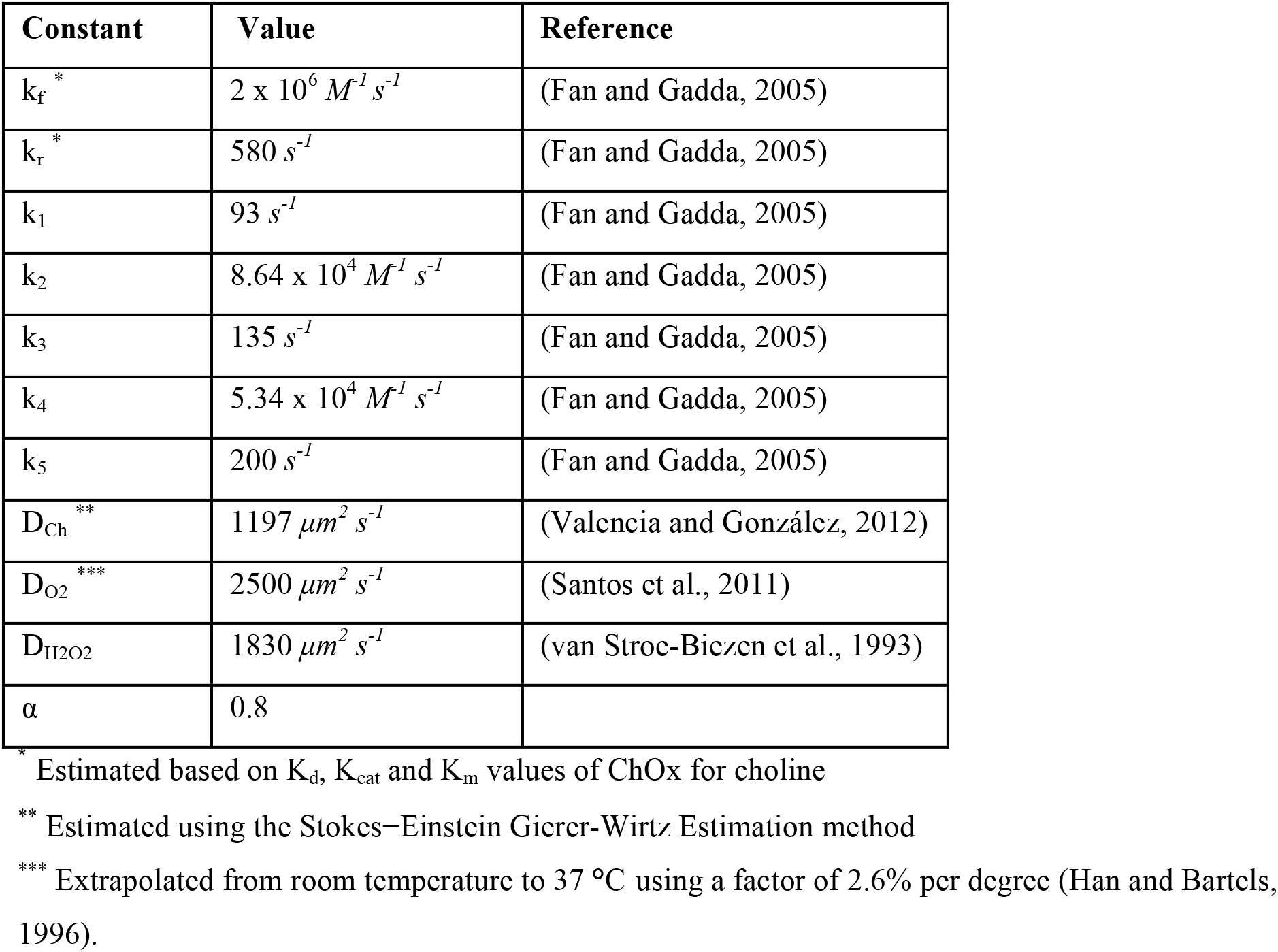
Values of constants used in the biosensor model

| Constant | Value | Reference |
| --- | --- | --- |
| $k_f^*$ | $2 \times 10^6 M^{-1} s^{-1}$ | (Fan and Gadda, 2005) |
| $k_r^*$ | $580 s^{-1}$ | (Fan and Gadda, 2005) |
| $k_1$ | $93 s^{-1}$ | (Fan and Gadda, 2005) |
| $k_2$ | $8.64 \times 10^4 M^{-1} s^{-1}$ | (Fan and Gadda, 2005) |
| $k_3$ | $135 s^{-1}$ | (Fan and Gadda, 2005) |
| $k_4$ | $5.34 \times 10^4 M^{-1} s^{-1}$ | (Fan and Gadda, 2005) |
| $k_5$ | $200 s^{-1}$ | (Fan and Gadda, 2005) |
| $D_{Ch}^{**}$ | $1197 \mu m^2 s^{-1}$ | (Valencia and González, 2012) |
| $D_{O_2}^{***}$ | $2500 \mu m^2 s^{-1}$ | (Santos et al., 2011) |
| $D_{H_2O_2}$ | $1830 \mu m^2 s^{-1}$ | (van Stroe-Biezen et al., 1993) |
| $\alpha$ | 0.8 | |

\* Estimated based on $K_d$ , $K_{cat}$ and $K_m$ values of ChOx for choline
\*\* Estimated using the Stokes–Einstein Gierer-Wirtz Estimation method
\*\*\* Extrapolated from room temperature to 37 °C using a factor of 2.6% per degree (Han and Bartels, 1996).

#### Statistical analysis

All statistical tests were performed using Matlab. Prior to statistical comparisons, the normality of the data was tested by a Anderson-Darling test. In the case of non-normal distributions, comparisons between two groups were performed using a non-parametric two-sided sign test (signtest, Matlab). To test the effect of two factors on non-normal data (effect of ripple count and power on COA or O_2_ transients), an aligned-rank-transformation (Wobbrock et al., 2011) was applied followed by two-way ANOVA for unbalanced data (nanova, Matlab). Whenever normality could not be discarded, twosided two-sample *t*-tests (paired or unpaired) were used to compare two groups. Two-sided one-sample *t*-tests were used to test deviations from the null hypothesis (zero). Multiple comparisons were done by one-way ANOVA for unbalanced data. The effect of two or more factors was accounted for by two- or three-way ANOVA for unbalanced data.

Data were presented as average ± 95% confidence interval (CI) to allow easy assessment of the significance of estimates. Average corresponds to the mean in normal data or to the median otherwise. The CI of medians was computed based on fractional order statistics (Hutson, 1999). Confidence intervals for correlation coefficients and COA vs. O_2_ slopes were computed from the percentiles 2.5 and 97.5 of bootstrapped data.

## Acknowledgments

This work was supported by Bundesministerium für Bildung und Forschung [grant number 01GQ0440] and the Munich Cluster for Systems Neurology [grant number SyNergy EXC 1010]. We would like to thank Francisco Almeida and Kenneth Klau for technical assistence.

## Competing interests

The authors declare no competing interests.

## Supplementary Materials

**Figure S1.**
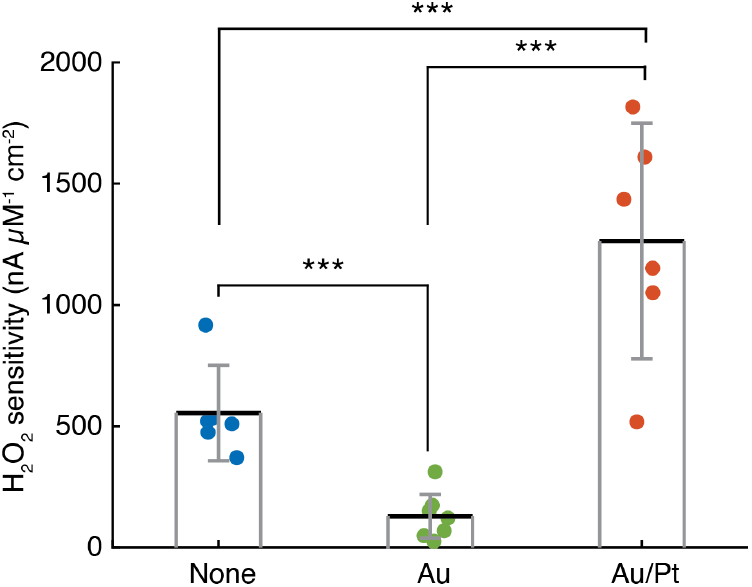
Effect of platings on platinum electrode sensitivity to H_2_O_2_. Normalized H_2_O_2_ sensitivity of Pt/Ir disc electrodes (n=6, data from our previous sensor design (Santos et al., 2015)), gold-plated Pt/Ir tetrode sites (n=7) and gold/platinum plated tetrode sites (n=6). All electrode surfaces were coated with chitosan/ChOx matrices. All group pairwise comparisons were statistically significant (p<0.001, one-way ANOVA for unbalanced data followed by Tukey-Kramer post-hoc test). *** p<0.001

**Figure S2.**
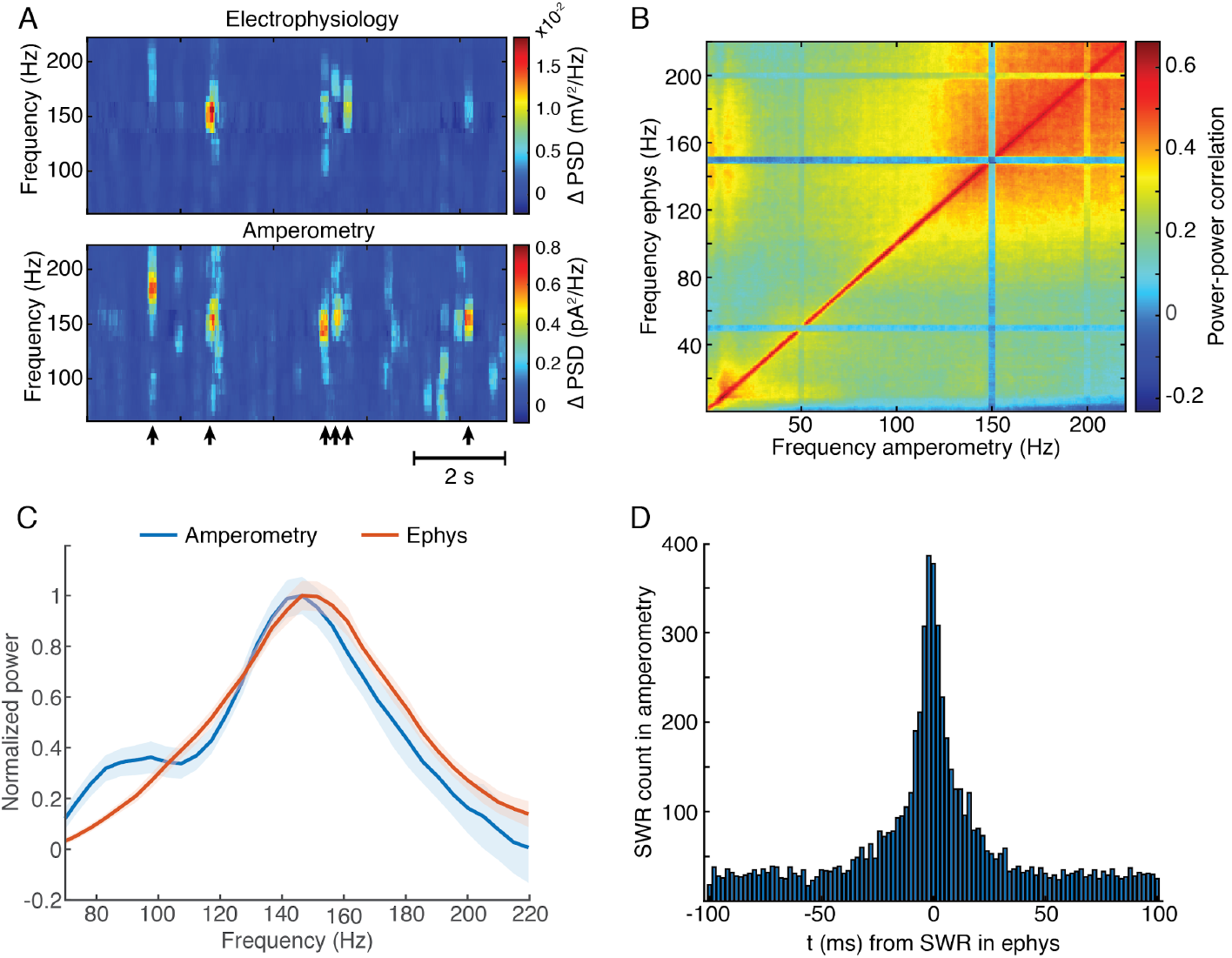
Amperometric currents reliably track LFP spectral content over a wide frequency range. (A) Representative power spectrogram from high-frequency range of electrophysiology and amperometry derived LFP signals in the CA1 pyramidal layer during NREM sleep. Arrows indicate timings of SWRs detected from LFP. (B) Comodugram showing the correlation between LFP power, recorded during NREM sleep, from a silicon probe’s site in CA1 pyramidal layer and the power of the amperometric signal from a biosensor site, targeted to the same hippocampal layer. (C) Normalized power spectra triggered to SWRs using both modalities. (D) Cross-correlogram of SWR timings detected using amperometry and electrophysiology (n=8074 for electrophysiology and n=8085 for amperometry).

**Figure S3.**
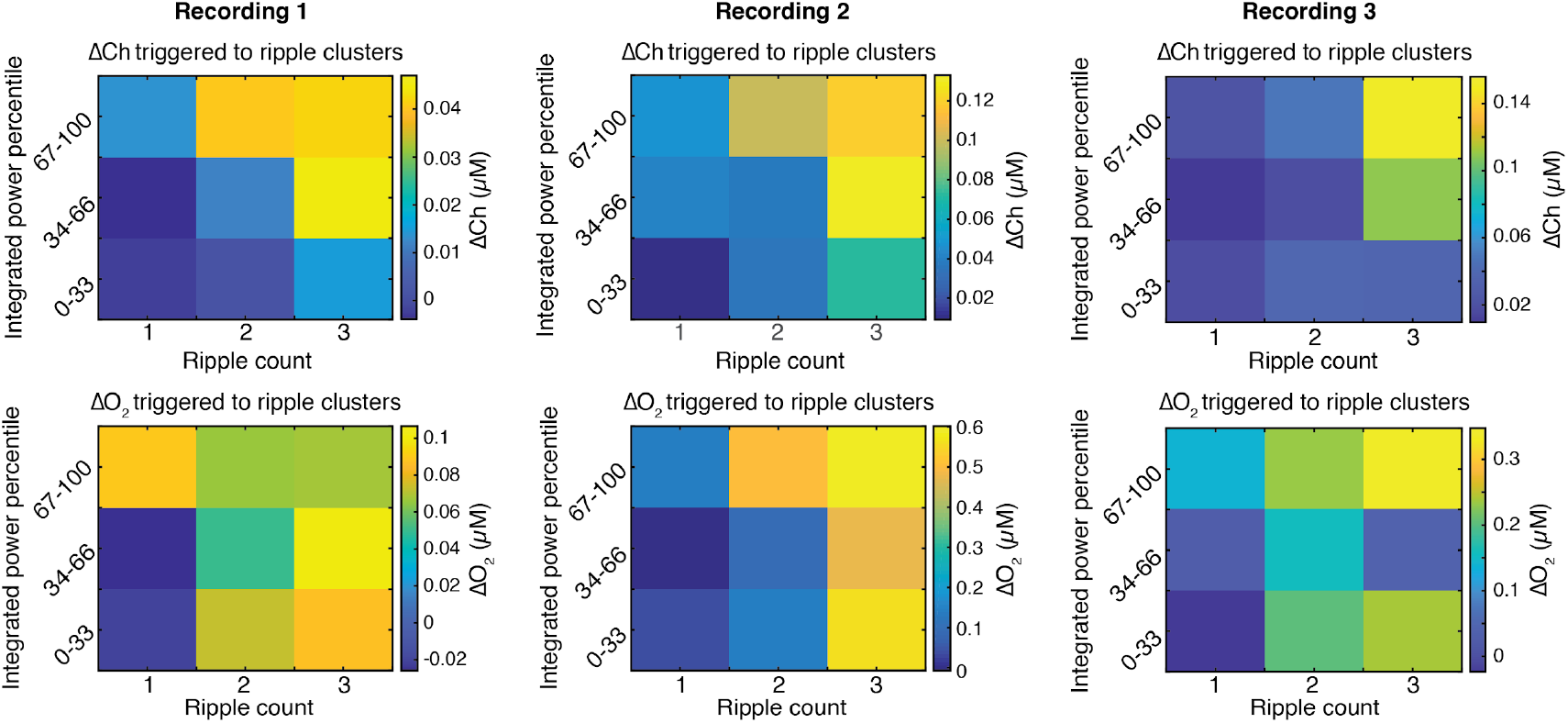
Phasic COA and oxygen responses following SWRs jointly depend on their power and grouping. Amplitude of ChOx activity and O_2_ transients as a function of SWR count in a 2 s time window, sorted by different percentile ranges of summed ripple power. The data was collected from three recording sessions. Statistic tests were performed by two-way ANOVA for unbalanced data following ART to account for non-gaussian distributions. In recording session 1, both SWR count and total ripple power significantly affected the amplitude of ChOx and O_2_ (*p*<0.005 and *F*>5.5 for both factors in ChOx and O_2_ data). The same applied for recording 2 (*p*<0.0001 and *F*>12 for both factors in ChOx and O_2_ data). As for recording session 3, the SWR count effect was significant for both sensor signals (p<0.05 and F>3.4) and the summed ripple power significantly affected O_2_ (p<0.05 and F>4), but was not consistently related to ChOx transients amplitude (p=0.07, F=2.71).

**Figure S4.**
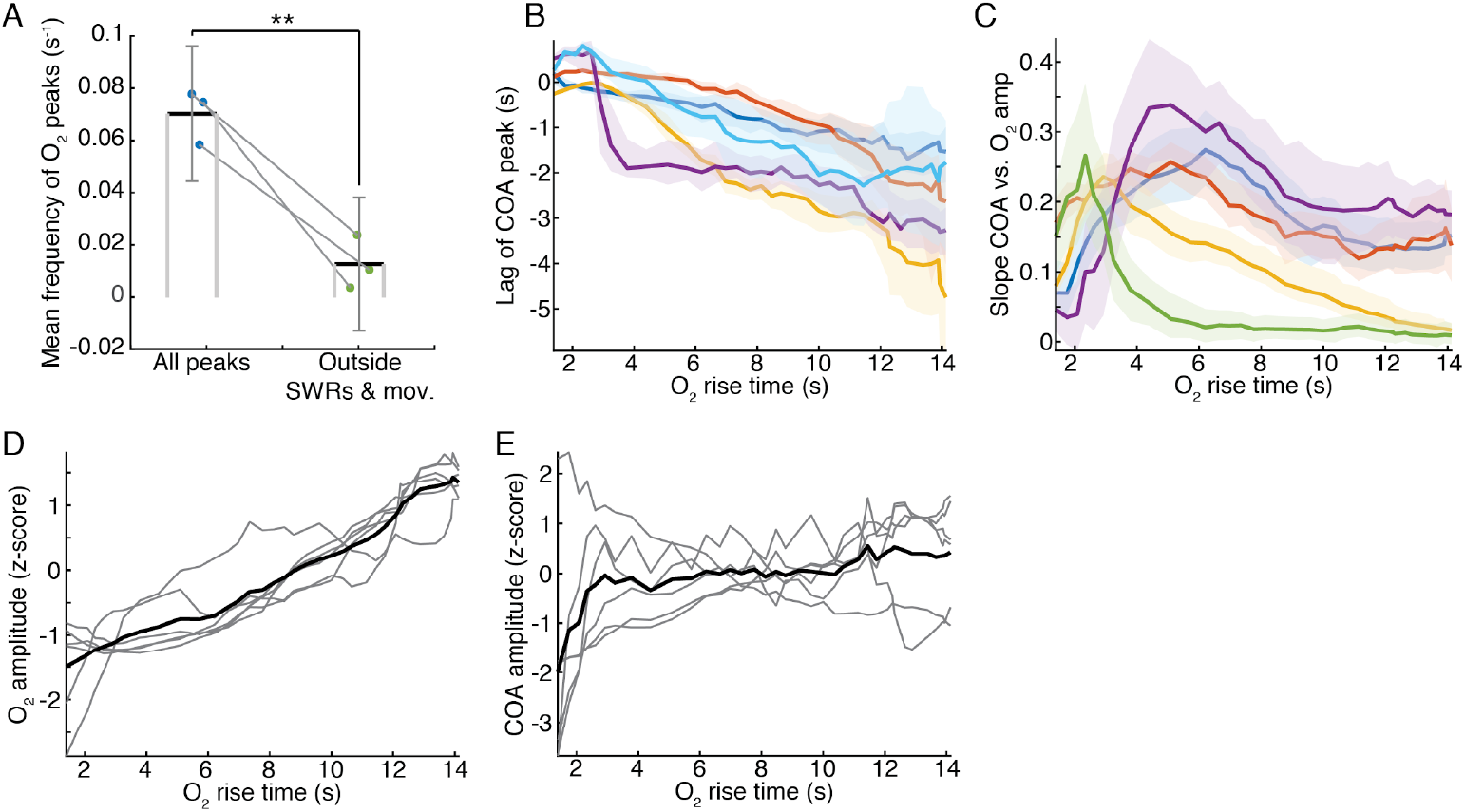
COA dynamics associated with O_2_ transients across all experiments. (A) Mean occurrence rate of O_2_ peaks during either whole recordings or periods outside SWRs and locomotion bouts (events occurring within −5 s to +1 s from SWRs or within −14 s to + 4 s from peaks in speed were excluded). The difference between O_2_ peak rates was significant (p<0.01, paired *t*-test). (B-C) plots display the same statistics as in an example on Figure 5E, but for all recordings (B) Lag of ChOx activity peaks relative to O_2_ for changes in O_2_ detected with varying rise time (from whole recordings, n=5). Colors represent data from different recording sessions shown as medians ± CIs. (C) Slope of COA response *vs.* O_2_ amplitude for O_2_ transients detected with a varying rise time. Colors represent data from different recording sessions shown as medians ± CIs. (D) z-scored amplitude of detected O_2_ peaks as a function of O_2_ rise time. Each trace represents medians from each recording. (E) z-scored amplitude of COA transients peaks associated with O_2_ transients as a function of O_2_ rise time. Each trace represents medians from each recording.

**Figure S5.**
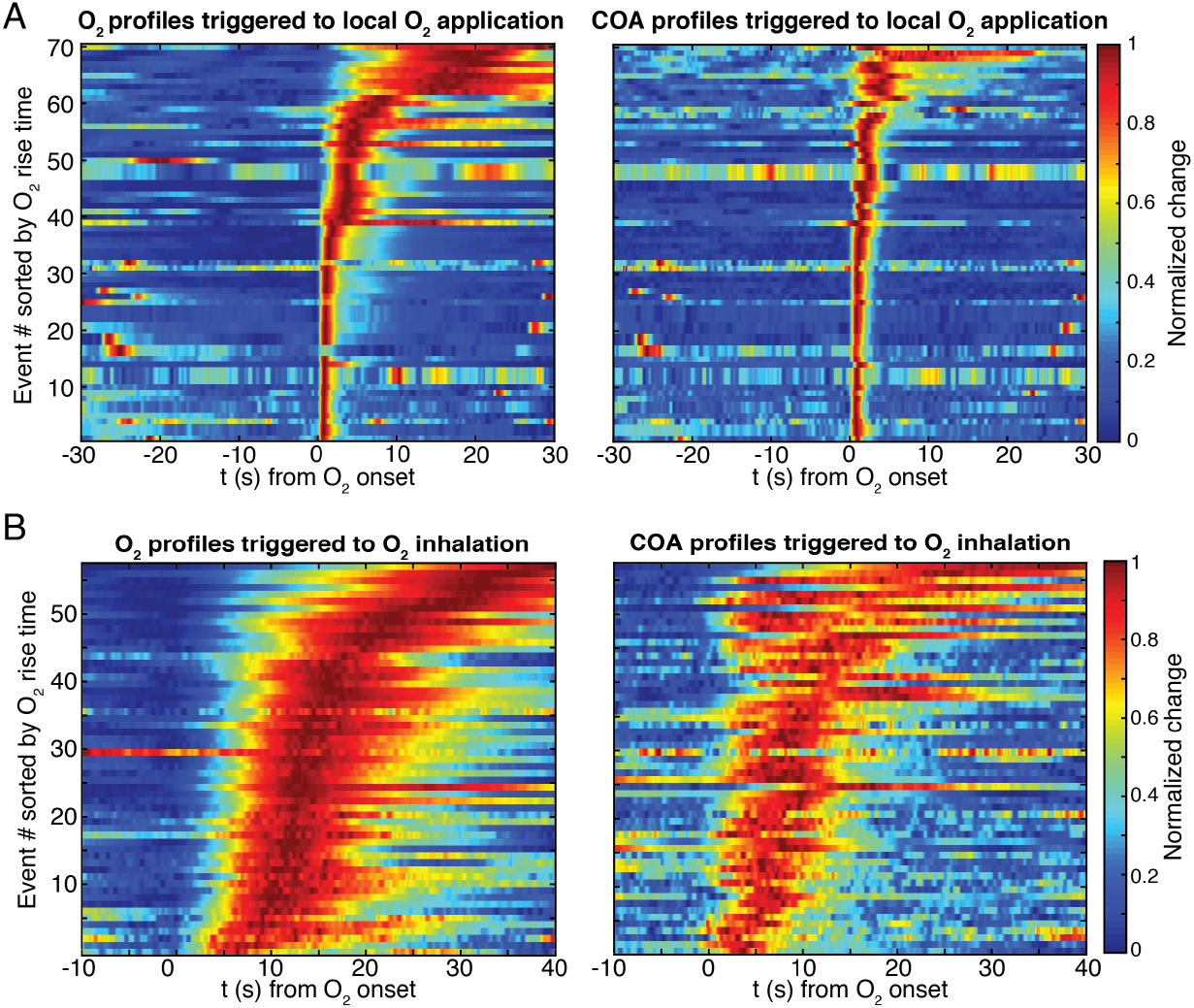
Raw sensor responses evoked by exogenous oxygen *in vivo.* (A) Normalized O_2_ and COA transients evoked by local O_2_ application, sorted by O_2_ rise time. Events were obtained from three recording sessions (n=13-40 per session). (B) Normalized O_2_ and COA transients evoked by O_2_ inhalation, sorted by O_2_ rise time. Events were obtained from three recording sessions (n=8-40 per session).

**Figure S6.**
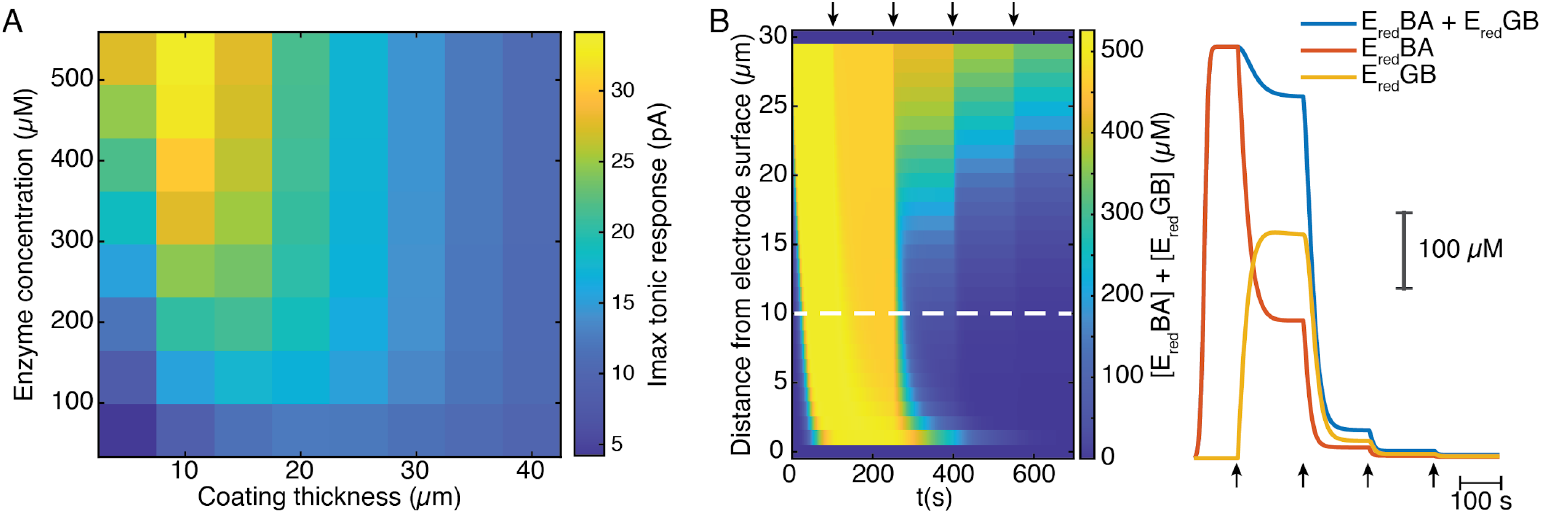
Mathematical simulation of ChOx-biosensor tonic responses and reduced enzymebound intermediate profiles. (A) Maximal cumulative tonic sensor responses as a function of coating thickness and enzyme concentration in the coating. (B) Left shows simulated concentration profile of total reduced enzyme-bound intermediates (E_red_BA+E_red_GB) in the sensor coating as a function of distance from the electrode surface during a simulated calibration of a high enzyme-loaded sensor (coating thickness of 30 μm and enzyme concentration of 560 μM). Right shows the concentration profile of total and individual enzyme-bound intermediates at 10 μm from the electrode surface (dashed line in left panel). Arrows indicate 5 μM O_2_ step increments in solution.

## Notes

### Competing Interest Statement

The authors have declared no competing interest.

## References

Baker, K.L., Bolger, F.B., and Lowry, J.P. (2015). A microelectrochemical biosensor for real-time in vivo monitoring of brain extracellular choline. Anal. 140, 3738–3745.

Baker, K.L., Bolger, F.B., and Lowry, J.P. (2017). Development of a microelectrochemical biosensor for the real-time detection of choline. Sensors Actuators B: Chem. 243.

Baronas, R., Ivanauskas, F., and Kulys, J. (2009). Mathematical modeling of biosensors: an introduction for chemists and mathematicians.

Brehm, R., Lindmar, R., and Löffelholz, K. (1987). Muscarinic mobilization of choline in rat brain in vivo as shown by the cerebral arterio-venous difference of choline. J. Neurochem. 48, 1480–1485.

Burke, L.D. (1996). Anomalous Oxidation Reactions at Noble Metal Surfaces at Low Potentials: With Particular Reference to Palladium. J. Electrochem. Soc. 143.

Burke, L.D., and Nugent, P.F. (1997). The electrochemistry of gold: I the redox behaviour of the metal in aqueous media. Gold Bull. 30.

Burmeister, J.J., Palmer, M., and Gerhardt, G.A. (2003). Ceramic-based multisite microelectrode array for rapid choline measures in brain tissue. Anal. Chim. Acta 481.

Buzsáki, G. (2002). Theta oscillations in the hippocampus. Neuron 33, 325–340.

Buzsáki, G. (2015). Hippocampal sharp wave-ripple: A cognitive biomarker for episodic memory and planning. Hippocampus 25, 1073–1188.

Chatard, C., Meiller, A., and Marinesco, S. (2018). Microelectrode Biosensors for in vivo Analysis of Brain Interstitial Fluid. Electroanalysis 30.

Csernai, M., Borbély, S., Kocsis, K., Burka, D., Fekete, Z., Balogh, V., Káli, S., Emri, Z., and Barthó, P. (2019). Dynamics of sleep oscillations is coupled to brain temperature on multiple scales. J. Physiol.

Dixon, B.M., Lowry, J.P., and O’Neill, R.D. (2002). Characterization in vitro and in vivo of the oxygen dependence of an enzyme/polymer biosensor for monitoring brain glucose. J. Neurosci. Methods 119, 135–142.

Domínguez-Domínguez, S., Arias-Pardilla, J., Berenguer-Murcia, Á., Morallón, E., and Cazorla-Amorós, D. (2008). Electrochemical deposition of platinum nanoparticles on different carbon supports and conducting polymers. J. Appl. Electrochem. 38.

Eggermann, E., Kremer, Y., Crochet, S., and Petersen, C.C.H. (2014). Cholinergic signals in mouse barrel cortex during active whisker sensing. Cell Reports 9, 1654–1660.

Fan, F., and Gadda, G. (2005). On the catalytic mechanism of choline oxidase. J. Am. Chem. Soc. 127, 2067–2074.

Ferguson, J.E., Boldt, C., and Redish, A.D. (2009). Creating low-impedance tetrodes by electroplating with additives. Sensors Actuators. Phys. 156, 388–393.

Garguilo, M.G., and Michael, A.C. (1996). Amperometric microsensors for monitoring choline in the extracellular fluid of brain. J. Neurosci. Methods 70, 73–82.

Gray, C.M., Maldonado, P.E., Wilson, M., and McNaughton, B. (1995). Tetrodes markedly improve the reliability and yield of multiple single-unit isolation from multi-unit recordings in cat striate cortex. J. Neurosci. Methods 63, 43–54.

Gu, Z., Alexander, G.M., Dudek, S.M., and Yakel, J.L. (2017). Hippocampus and Entorhinal Cortex Recruit Cholinergic and NMDA Receptors Separately to Generate Hippocampal Theta Oscillations. Cell Reports 21, 3585–3595.

Han, P., and Bartels, D.M. (1996). Temperature Dependence of Oxygen Diffusion in H2O and D2O. The Journal of Physical Chemistry 100.

Hangya, B., Ranade, S.P., Lorenc, M., and Kepecs, A. (2015). Central Cholinergic Neurons Are Rapidly Recruited by Reinforcement Feedback. Cell 162, 1155–1168.

Hascup, K.N., Hascup, E.R., Littrell, O.M., Hinzman, J.M., Werner, C.E., Davis, V.A., Burmeister, J.J., Pomerleau, F., Quintero, J.E., Huettl, P., et al. (2013). Microelectrode Array Fabrication and Optimization for Selective Neurochemical Detection. In Microelectrode Biosensors, (Humana Press), pp. 27–54.

Hashemi, P., Walsh, P.L., Guillot, T.S., Gras-Najjar, J., Takmakov, P., Crews, F.T., and Wightman, R.M. (2011). Chronically Implanted, Nafion-Coated Ag/AgCl Reference Electrodes for Neurochemical Applications. ACS Chem. Neurosci. 2, 658–666.

Hasselmo, M.E., and McGaughy, J. High acetylcholine levels set circuit dynamics for attention and encoding and low acetylcholine levels set dynamics for consolidation. Prog Brain Res 145, 207–231.

Hasselmo, M.E., and McGaughy, J. (2004). High acetylcholine levels set circuit dynamics for attention and encoding and low acetylcholine levels set dynamics for consolidation. Prog. Brain Res. 145, 207–231.

Hekmat, A., Saboury, A.A., Moosavi-Movahedi, A.A., Ghourchian, H., and Ahmad, F. (2008). Effects of pH on the activity and structure of choline oxidase from Alcaligenes species. Acta Biochim. Pol. 55, 549–557.

Howe, W.M., Gritton, H.J., Lusk, N.A., Roberts, E.A., Hetrick, V.L., Berke, J.D., and Sarter, M. (2017). Acetylcholine Release in Prefrontal Cortex Promotes Gamma Oscillations and Theta-Gamma Coupling during Cue Detection. J. Neurosci. Off. J. Soc. Neurosci. 37, 3215–3230.

Hutson, A.D. (1999). Calculating nonparametric confidence intervals for quantiles using fractional order statistics. J. Appl. Stat. 26, 343–353.

Lamers-Lemmers, J.P., Hoofd, L.J., and Oeseburg, B. (2000). Non-steady-state O(2) diffusion in metmyoglobin solutions studied in a diffusion chamber. Biochem. Biophys. Res. Commun. 276, 773–778.

Leithner, C., and Royl, G. (2014). The oxygen paradox of neurovascular coupling. J. Cereb. Blood Flow Metab. Off. J. Int. Soc. Cereb. Blood Flow Metab. 34, 19–29.

Li, S., Topchiy, I., and Kocsis, B. (2007). The effect of atropine administered in the medial septum or hippocampus on high-and low-frequency theta rhythms in the hippocampus of urethane anesthetized rats. Synap. 61, 412–419.

Lopes, G., Bonacchi, N., Frazão, J., Neto, J.P., Atallah, B.V., Soares, S., Moreira, L., Matias, S., Itskov, P.M., Correia, P.A., et al. (2015). Bonsai: an event-based framework for processing and controlling data streams. Front. Neuroinformatics 9, 7.

Lovett-Barron, M., Kaifosh, P., Kheirbek, M.A., Danielson, N., Zaremba, J.D., Reardon, T.R., Turi, G.F., Hen, R., Zemelman, B.V., and Losonczy, A. (2014). Dendritic inhibition in the hippocampus supports fear learning. Sci. 343, 857–863.

Lyons, D.G., Parpaleix, A., Roche, M., and Charpak, S. (2016). Mapping oxygen concentration in the awake mouse brain. ELife 5.

Marrosu, F., Portas, C., Mascia, M.S., Casu, M.A., Fà, M., Giagheddu, M., Imperato, A., and Gessa, G.L. (1995). Microdialysis measurement of cortical and hippocampal acetylcholine release during sleep-wake cycle in freely moving cats. Brain Res. 671, 329–332.

McMahon, C.P., Rocchitta, G., Kirwan, S.M., Killoran, S.J., Serra, P.A., Lowry, J.P., and O’Neill, R.D. (2007). Oxygen tolerance of an implantable polymer/enzyme composite glutamate biosensor displaying polycation-enhanced substrate sensitivity. Biosens. & Bioelectron. 22, 1466–1473.

Mikulovic, S., Restrepo, C.E., Siwani, S., Bauer, P., Pupe, S., Tort, A.B.L., Kullander, K., and Leão, R.N. (2018). Ventral hippocampal OLM cells control type 2 theta oscillations and response to predator odor. Nat. Commun. 9, 3638.

Mitra, P.P., and Pesaran, B. (1999). Analysis of dynamic brain imaging data. Biophys. J. 76, 691–708.

Murr, R., Berger, S., Schürer, L., Peter, K., and Baethmann, A. (1994). A novel, remote-controlled suspension device for brain tissue PO2 measurements with multiwire surface electrodes. Pflugers Arch. Eur. J. Physiol. 426, 348–350.

Nair, P.K., Buerk, D.G., and Halsey, J.H. (1987). Comparisons of oxygen metabolism and tissue PO2 in cortex and hippocampus of gerbil brain. Stroke 18, 616–622.

Njagi, J., Ispas, C., and Andreescu, S. (2008). Mixed ceria-based metal oxides biosensor for operation in oxygen restrictive environments. Anal. Chem. 80, 7266–7274.

Norimoto, H., Mizunuma, M., Ishikawa, D., Matsuki, N., and Ikegaya, Y. (2012). Muscarinic receptor activation disrupts hippocampal sharp wave-ripples. Brain Res. 1461, 1–9.

O’Neill, R.D., Chang, S.-C., Lowry, J.P., and McNeil, C.J. (2004). Comparisons of platinum, gold, palladium and glassy carbon as electrode materials in the design of biosensors for glutamate. Biosens. & Bioelectron. 19, 1521–1528.

Parikh, V., Pomerleau, F., Huettl, P., Gerhardt, G. a, Sarter, M., and Bruno, J.P. (2004). Rapid assessment of in vivo cholinergic transmission by amperometric detection of changes in extracellular choline levels. Eur. J. Neurosci. 20, 1545–1554.

Parikh, V., Kozak, R., Martinez, V., and Sarter, M. (2007). Prefrontal acetylcholine release controls cue detection on multiple timescales. Neuron 56, 141–154.

Park, J., Kim, C.-S., and Choi, M. (2006). Oxidase-Coupled Amperometric Glucose and Lactate Sensors With Integrated Electrochemical Actuation System. IEEE Trans. Instrum. Meas. 55.

Ramirez-Villegas, J.F., Logothetis, N.K., and Besserve, M. (2015). Diversity of sharp-wave-ripple LFP signatures reveals differentiated brain-wide dynamical events. Proc. Natl. Acad. Sci. United States Am. 112, E6379–E6387.

Reimer, J., McGinley, M.J., Liu, Y., Rodenkirch, C., Wang, Q., McCormick, D.A., and Tolias, A.S. (2016). Pupil fluctuations track rapid changes in adrenergic and cholinergic activity in cortex. Nat. Commun. 7, 13289.

Santos, R.M., Lourenço, C.F., Piedade, A.P., Andrews, R., Pomerleau, F., Huettl, P., Gerhardt, G.A., Laranjinha, J., and Barbosa, R.M. (2008). A comparative study of carbon fiber-based microelectrodes for the measurement of nitric oxide in brain tissue. Biosens. & Bioelectron. 24, 704–709.

Santos, R.M., Lourenço, C.F., Gerhardt, G.A., Cadenas, E., Laranjinha, J., and Barbosa, R.M. (2011). Evidence for a pathway that facilitates nitric oxide diffusion in the brain. Neurochem. Int. 59, 90–96.

Santos, R.M., Laranjinha, J., Barbosa, R.M., and Sirota, A. (2015). Simultaneous measurement of cholinergic tone and neuronal network dynamics in vivo in the rat brain using a novel choline oxidase based electrochemical biosensor. Biosens. & Bioelectron. 69, 83–94.

Siegle, J.H., López, A.C., Patel, Y.A., Abramov, K., Ohayon, S., and Voigts, J. (2017). Open Ephys: an open-source, plugin-based platform for multichannel electrophysiology. J. Neural Eng. 14, 045003.

Steriade, M. (2004). Acetylcholine systems and rhythmic activities during the waking-sleep cycle. Prog. Brain Res. 145, 179–196.

van Stroe-Biezen, S.A.., Everaerts, F.., Janssen, L.J.., and Tacken, R.. (1993). Diffusion coefficients of oxygen, hydrogen peroxide and glucose in a hydrogel. Anal. Chim. Acta 273.

Takata, N., Nagai, T., Ozawa, K., Oe, Y., Mikoshiba, K., and Hirase, H. (2013). Cerebral blood flow modulation by Basal forebrain or whisker stimulation can occur independently of large cytosolic Ca2+ signaling in astrocytes. PloS One 8, e66525.

Teles-Grilo Ruivo, L.M., and Mellor, J.R. (2013). Cholinergic modulation of hippocampal network function. Front. Synaptic Neurosci. 5, 2.

Teles-Grilo Ruivo, L.M., Baker, K.L., Conway, M.W., Kinsley, P.J., Gilmour, G., Phillips, K.G., Isaac, J.T.R., Lowry, J.P., and Mellor, J.R. (2017). Coordinated Acetylcholine Release in Prefrontal Cortex and Hippocampus Is Associated with Arousal and Reward on Distinct Timescales. Cell Reports 18, 905–917.

Valencia, D.P., and González, F.J. (2012). Estimation of diffusion coefficients by using a linear correlation between the diffusion coefficient and molecular weight. Journal of Electroanalytical Chemistry 681.

Vandecasteele, M. M. S., Royer, S., Belluscio, M., Berényi, A., Diba, K., Fujisawa, S., Grosmark, A., Mao, D., Mizuseki, K., et al. (2012). Large-scale recording of neurons by movable silicon probes in behaving rodents. J. Vis. Exp. JoVE e3568.

Vandecasteele, M., Varga, V., Berényi, A., Papp, E., Barthó, P., Venance, L., Freund, T.F., and Buzsáki, G. (2014). Optogenetic activation of septal cholinergic neurons suppresses sharp wave ripples and enhances theta oscillations in the hippocampus. Proc. Natl. Acad. Sci. United States Am. 111, 13535–13540.

Venton, B.J., Michael, D.J., and Wightman, R.M. (2003). Correlation of local changes in extracellular oxygen and pH that accompany dopaminergic terminal activity in the rat caudate-putamen. J. Neurochem. 84, 373–381.

Viggiano, A., Marinesco, S., Pain, F., Meiller, A., and Gurden, H. (2012). Reconstruction of field excitatory post-synaptic potentials in the dentate gyrus from amperometric biosensor signals. J. Neurosci. Methods 206, 1–6.

Wobbrock, J.O., Findlater, L., Gergle, D., and Higgins, J.J. (2011). The aligned rank transform for nonparametric factorial analyses using only anova procedures. In Proceedings of the 2011 Annual Conference on Human Factors in Computing Systems-CHI, 11, (ACM Press), p. 143.

Zhang, H., Lin, S.-C., and Nicolelis, M.A.L. (2009). Acquiring local field potential information from amperometric neurochemical recordings. J. Neurosci. Methods 179, 191–200.

Zhang, H., Lin, S.-C., and Nicolelis, M.A.L. (2010). Spatiotemporal coupling between hippocampal acetylcholine release and theta oscillations in vivo. J. Neurosci. Off. J. Soc. Neurosci. 30, 13431–13440.

Zhang, Q., Roche, M., Gheres, K.W., Chaigneau, E., Kedarasetti, T., Haselden, D., Charpak, S., Drew, P. J. (2019). Cerebral oxygenation during locomotion is modulated by respiration. Nat. Commun. 10, 5515.

